# Cell wall extensin arabinosylation is required for root directional response to salinity

**DOI:** 10.1101/2022.06.22.497042

**Authors:** Yutao Zou, Nora Gigli-Bisceglia, Eva van Zelm, Pinelopi Kokkinopoulou, Magdalena M. Julkowska, Maarten Besten, Thu-Phuong Nguyen, Hongfei Li, Jasper Lamers, Thijs de Zeeuw, Joram A. Dongus, Yuxiao Zeng, Yu Cheng, Iko T. Koevoets, Bodil Jørgensen, Marcel Giesbers, Jelmer Vroom, Tijs Ketelaar, Bent Larsen Petersen, Timo Engelsdorf, Joris Sprakel, Yanxia Zhang, Christa Testerink

## Abstract

Soil salinity is a major contributor to crop yield losses. To improve our understanding of root responses to salinity, we developed and exploit here a real-time salt-induced tilting assay (SITA). This method follows root growth upon both gravitropic and salt challenges, revealing that root bending upon tilting is modulated by salinity, but not by osmotic stress. Next, this salt-specific response was measured in 345 natural Arabidopsis accessions and we discovered a genetic locus, encoding for the cell-wall modifying enzyme EXTENSIN ARABINOSE DEFICIENT TRANSFERASE (ExAD), to be associated with root bending in salt. Extensins are a class of structural cell wall glycoproteins [hydroxyproline-rich glycoproteins (HRGPs)] which are post-translationally modified by O-glycosylation mostly in the form of hydroxyproline (Hyp)-arabinosylation. We show that salt induces ExAD-dependent Hyp-arabinosylation, influencing root bending responses and cell wall thickness. We report that roots of *exad* mutants, which lack extensin Hyp-Araf_4_ modifications, display increased root epidermal cell wall thickness and porosity and altered gravitropic root bending in salt, as well as a reduced salt avoidance response. Our results suggest that extensin modification via Hyp-arabinosylation represents a novel salt-specific cellular process that is required for the directional response of roots exposed to salinity.

## Introduction

Soil salinization is a major problem causing crop yield loss in agriculture, since most crops are sensitive to salt stress (Munns and Tester, 2008). Sodium chloride (NaCl) is the main factor causing soil salinity (FAO, 2021), affecting the plant on two levels: through osmotic stress, challenging the root water uptake, and through toxic accumulation of ions in various plant tissues (Hasegawa et al., 2000; Julkowska and Testerink, 2015). Excess of salt in soil thus obstructs various cellular processes and negatively impacts shoot and root growth (van Zelm et al., 2020). In the root, salt has been shown to modify root architecture and cell wall composition and to inhibit gravitropism [reviewed in (Zou et al., 2022)]. In response to salinity, cell wall modifications enable the increase of lignin and suberin deposition and in older tissues the formation of secondary cell walls, all contributing to limiting Na^+^ entry ((Voxeur et al., 2015; Karlova et al., 2021; Barberon et al., 2016; Duan et al., 2020). Moreover, salt treatment also modulates several general cell wall biosynthesis processes, such as cellulose synthesis and localization of cellulose microfibrils (Endler et al., 2015), galactan accumulation (Yan et al., 2021) and arabinose biosynthesis and metabolism (Zhao et al., 2019). In this study, we performed a genome wide association study (GWAS) of root growth parameters of Arabidopsis accessions acquired by employing a newly developed dynamic salt-induced tilting assay (SITA). Similar to the halotropic response (Deolu-Ajayi et al., 2019; Galvan-Ampudia et al., 2013), the salt-induced modulation of root direction in SITA is a Na^+^-specific response, that does not occur in response to osmotic stress or other monovalent cations, such as K^+^. We identified a region in the Arabidopsis genome that correlated with the ability of the root to change growth direction in response to salinity. Further investigation showed that the *locus* contains the EXTENSIN ARABINOSE DEFICIENT TRANSFERASE (ExAD) gene, encoding for an α-(1,3)-arabinosyltransferase that adds the fourth arabinose residue to the hydroxyproline of cell wall glycoproteins known as extensins. Extensins are hydroxyproline-rich glycoproteins (HRGPs) that are required for the primary cell wall architectural maintenance (Lamport et al., 2011; Liu et al., 2016). Crosslinking of extensins helps building up networks in the wall and contributes to the structural matrix (Cannon et al., 2008; Marzol et al., 2018). Extensins are post-translationally modified by hydroxylation and then subsequently by O-glycosylation, which includes serine-galactosylation and hydroxyproline (Hyp)-arabinosylation. In extensin Hyp-arabinosylation, several arabinosyltransferases have been identified to be involved in forming the Hyp-Araf_1-4_ side-chain at specific positions. HYDROXYPROLINE ARABINOSYLTRANSFERASES 1-3 (HPAT1-3) add the first arabinose (Hyp-Araf_1_) to hydroxyprolines (Hyps) (Ogawa-Ohnishi et al., 2013) and REDUCED RESIDUAL ARABINOSE 1-3 (RRA1-3), a β-(1,2)-arabinosyltransferase adds the second arabinose residue (Hyp-Araf_2_) (Velasquez et al., 2011; Egelund et al., 2007). Next, XYLOGLUCAN ENDOGLUCANASE 113 (XEG113), also a β-(1,2)-arabinosyltransferase, adds the third arabinose residue (Hyp-Araf_3_) (Gille et al., 2009), and finally ExAD adds the fourth residue (Hyp-Araf_4_) (Møller et al., 2017). Recent studies suggest that extensin crosslinking relies on the arabinosylation of extensin motifs, and that the cross-linked network may promote root defense to pathogen-derived elicitors and could limit colonization by pathogens (Castilleux et al., 2020). However, the role of arabinosylation in response to abiotic stress is yet unknown. Here, we reveal that ExAD is required for the salt-specific modulation of root direction and enhanced root epidermal cell wall thickness upon salt stress. We also display that the Hyp-Araf_4_-signal detected by a JIM11 antibody, (Castilleux et al., 2020; Xie et al., 2011; Pattathil et al., 2010; Smallwood et al., 1994) is ExAD dependent and increases in salt treated wilt type seedlings, suggesting that cell wall extensin arabinosylation is a cellular response to salinity. Together, our study shows that Hyp-Araf_4_ of cell wall proteins is involved in root cell wall modifications under salt stress conditions, and that ExAD is required for the modulation of root directional growth, revealing a crucial role for ExAD-mediated arabinosylation in root responses to salt stress.

## Results

### Root gravitropism is modified in a NaCl-specific manner

To understand the dynamics of root directional growth responses to salinity and investigate the sodium specificity of salt inhibition of gravitropism, different salts and osmotic treatments were applied in a salt-induced tilting assay (SITA) setup and dynamic root bending was recorded by a time-lapse imaging system. Four-day-old Arabidopsis Col-0 seedlings were transferred to ½ MS agar plates containing different treatments while plates were rotated 90 degrees clockwise to apply a gravistimulus, immediately followed by tracking of root growth and direction in a time-lapse system. Images were taken every 20 mins for at least 24 hours (Figure 1A upper panel). During image analysis, we quantified the Root Tip Direction (RTD) Angle as the angle of the vector diverging from the gravity vector, indicating root tip directionality, or root tip bending (Figure 1A lower panel). Root Tip Direction (RTD) was analyzed every 20 minutes from the application of salt and calculated only the direction of the root tip (on the last 10% of the total root length). This distinction is grounded in the observation that salt-treated roots exhibit altered gravitropic responses (Sun et al., 2008; Dinneny et al., 2008). Values within the interval of 0 to –90 degrees characterize the roots of control-treated seedlings, while angle intervals between 0 and 90 degrees are typical of salt-stressed roots. Consistently with previous studies (Sun et al., 2008; Dinneny et al., 2008) the analysis performed in the SITA set-up displayed altered root tip direction consistent with an altered root gravitropic response. Changes in root bending through RTD were detected already during early time points (Figure 1B). Interestingly, while 100 mM NaCl treatment significantly affected root tip direction (angle intervals between 0 and 90 degrees) the RTDs of seedlings transplanted to 200 mM sorbitol or 100 mM KCl were more similar to those quantified in control condition (Figure 1B; Supplemental Video), even though NaCl, KCl and sorbitol treatments all resulted in a reduced root growth rate when compared to control conditions (Figure 1C) indicating that the difference in RTD is specific to NaCl and not likely caused by growth rate inhibition. NaNO_3_ treatment had a similar effect as NaCl, confirming that it is the presence of Na^+^ ions that triggers the altered root directionality in SITA (Figure S1A). To assess a possible effect of root developmental stages, we repeated the experiments by using 3-day-old Col-0 seedlings, and similar results were observed (Figure S1B, C). We also tested the effects on plant response of a 90-degree re-orientation of the plates towards either right (clockwise) or left (anticlockwise) direction. 90-degree re-orientation of the plates, shows that after the initial phase, NaCl specifically modified root directional growth in both cases at later stage to a consistent right re-orientation angle (Figure S1D). Furthermore, we found a dose-dependent effect on the salt-dependent changes in root direction when testing a concentration range from 0 mM, 75 mM, 100 mM, and 125 mM in SITA (Figure 1D, E), and we selected a concentration of 100 mM NaCl treatment for a further natural diversity panel screen.

**Figure 1.**
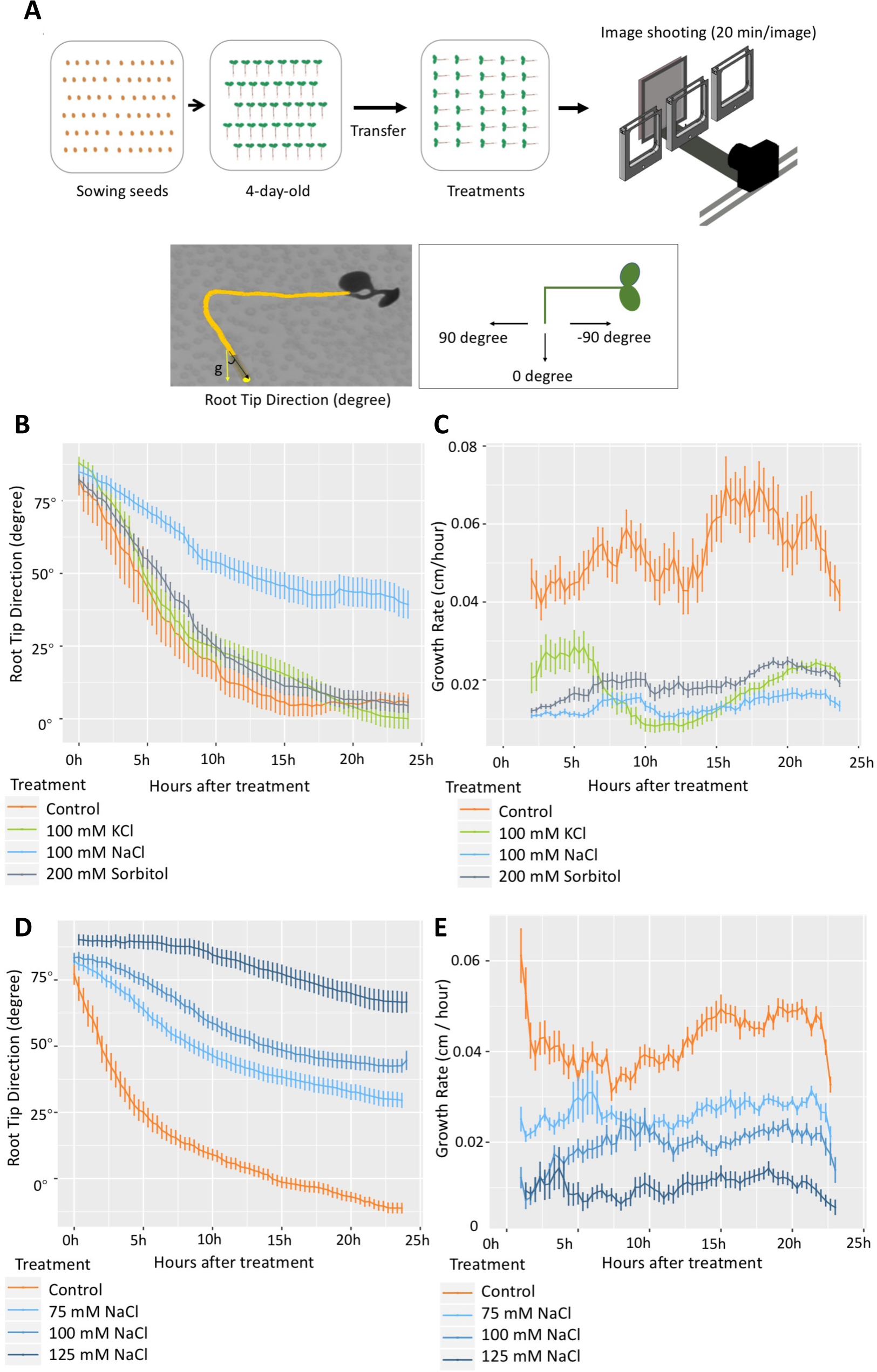
Root Tip direction (RTD) in time-lapse SITA is specifically modulated by NaCl treatment in a dose-dependent manner. **A**, Experimental set-up of SITA. Four-day-old seedlings germinated on ½ MS medium were transferred to ½ MS agar plates containing different treatments, and at the same time plates were rotated 90 degrees anticlockwise. The root dynamic growth response was traced in a time-lapse system with an imaging frequency of one image per 20 min per plate. **B**, Root tip direction (RTD) values expressed as degree angle (Y-axis) were quantified every 20 min in time lapse set-up in 4-day-old Col-0 seedlings under different treatments (control, 100 mM NaCl, 100 mM KCl and 200 mM sorbitol). **C**, Growth rate of seedlings roots of experiment B expressed in cm/h was analyzed over 24 h treatment. **D**, RTD values quantified in 4-day old treated Col-0 seedlings under different concentrations of NaCl (0 mM, 75 mM, 100 mM and 125 mM) over 24 h. **E**, Shows the growth rates of roots treated as in D. RTD angle values and root growth rates were obtained with SmartRoot. Values represent means ± SE from 30 seedlings, error bars represent SE of the mean. Data are representative of 3 independent experiments.

### GWAS analysis on Arabidopsis natural accessions through SITA revealed candidate loci associated with alteration in root directionality

Given the salt-specificity of the SITA response, we used it as a phenotypic readout to identify new molecular players that alter the salt-dependent root bending response during gravistimulation. SITA root growth parameters were collected from 345 Arabidopsis HapMap accessions (Table S1). From an initial survey performed on 20 Arabidopsis accessions, the overall root tip directional growth response observed in these accessions revealed a multi-phasic, response pattern (Figure S2), characterized by 3 distinct phases (Phase I, II and III) during the 30 h of treatment. Phase I (0 h –10 h) describes the acceleration stage in which the RTD of the tested accessions starts to respond to gravitropism. For most accessions, a strong effect of 100 mM NaCl on RTD was observed already during this phase, suggesting the inhibition of gravitropism by salt occurs immediately. During phase II (10 h – 20 h), most of the accessions exhibited a similar speed of change in root growth direction under both control and NaCl conditions. In Phase III (after 20 h), the root growth tends to be stabilized at a final direction after the initial effects of both gravitropism and salt stress (Figure S2). Because of the positive results obtained during this initial screen we expanded our analysis on the full HapMap diversity panel. The root phenotypic data obtained were subsequently analysed for correlation with SNP markers from the genome sequence of the Arabidopsis accessions (Alonso-Blanco et al., 2016) using a previously published GWAS R script, based on EMMAX and the ASReml R package (v.3.5.0) (Korte et al., 2012). In parallel the phenotypic data was analyzed with the online GWAPP web application (Seren et al., 2012). To map novel genetic loci associated with root bending response under salt application, we selected two different traits to be used as input for GWAS. The temporal traits K_RTD_^N/C^ (response in fitted rate of exponential decay) and root tip direction RTD^N-C^ (RTD under NaCl – RTD under Control) were analyzed at 5 h (Figure S3A, C, D and Table S3). The SNPs that were identified for the early response to salt stress are shown in Table S2 and S3. Together, the natural variation data and SNPs identified for these temporal traits provide a useful resource for future investigation of early responses of roots to salt. Next, we focused on the overall root bending response, which we expressed as Root Vector Angle (or RVA) as the angle of the vector, diverging from the gravity and following the direction of the root tip from the moment of the tilting upward until 23 h (Figure 2A). We measured RVA^N-C^ and observed that it was a highly consistent and robust trait displaying natural variation (Figure S3B). Hence, we employed the RVA^N-C^ at 23 h trait, to identify a genetic *locus* on chromosome 3 (Chr3) associated with 5 significant SNPs at or above the Bonferroni-corrected threshold, by using the GWAPP web tool (Figure 2B and Table S2). Consistently, when using the ASReml model (Korte et al., 2012) and a panel of 4,285,827 SNPs, also 5 SNPs above LOD 5.5 at the same positions were found associated with the RVA^N-C^ trait (Table S3, Figure S3B and S4). QQ plots indicate that the dataset of 345 Arabidopsis accessions for RVA^N-C^ at both 23 h traits exhibit conformity with a standard normal distribution.

**Figure 2.**
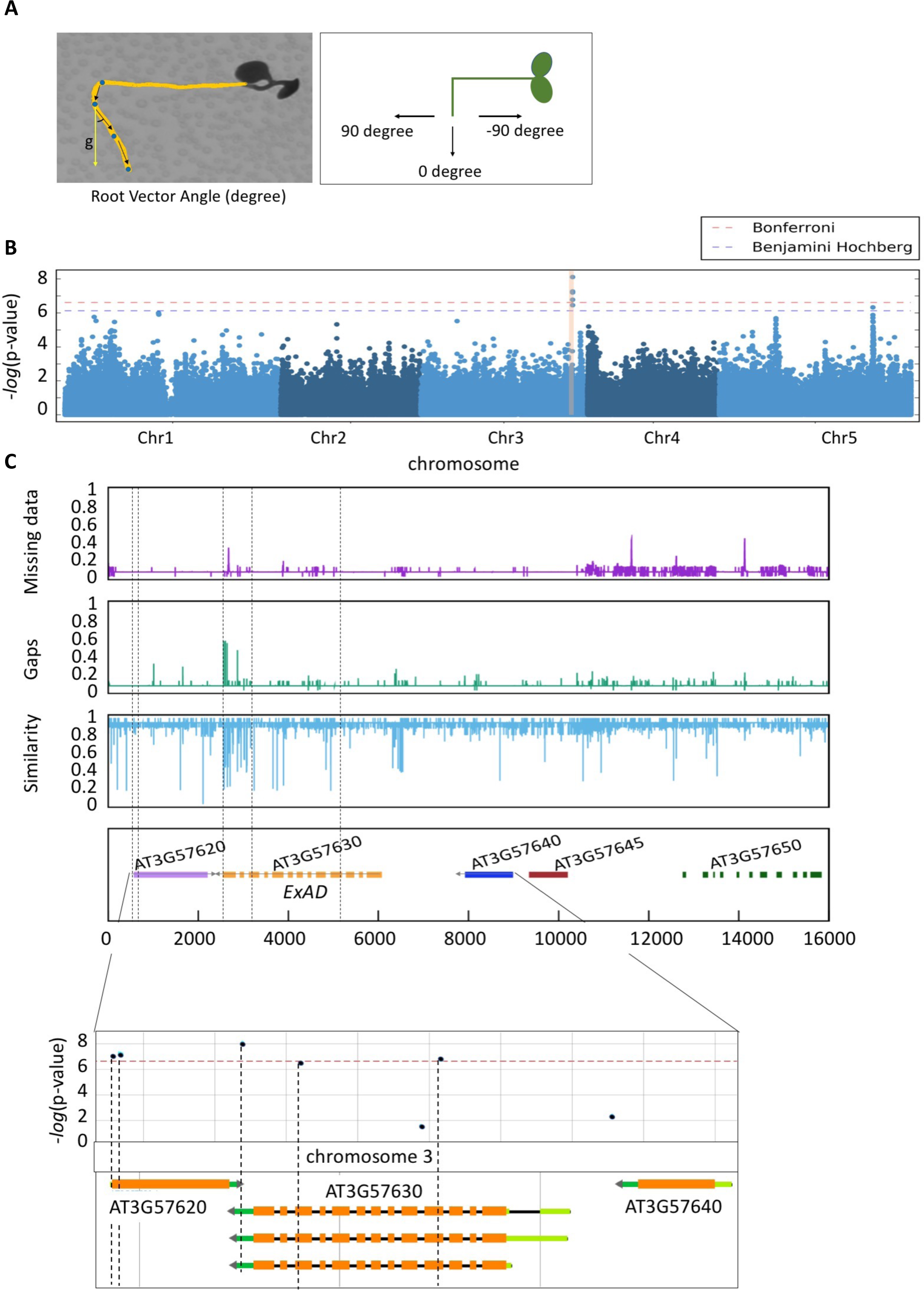
Natural variation in salt-modified root gravitropic response represented by root vector angle in SITA is associated with ExAD. **A**, Root vector angle (RVA) values, expressed as degree angle (Y-axis) were quantified every 23 h on daily scans. **B**, Manhattan plots for the SNPs associated with response root vector angle (RVA^N-C^) at 23 h under salt condition subtracted by RVA under control condition. In the phenotyping experiment for the GWAS, 5 replicates (seedlings) on each plate were used to calculate the average per accession, and Col-0 was repeated in each round of timelapse scanning as the standard control. QQ-plots are shown in Supplemental Figure S4. **C**, Divergence plot showing the genetic variation surrounding the significant SNPs on Chromosome 3 of all accessions in the 1001 genome database. The top purple graph represents the missing data, the middle green graph represents the gaps, and the bottom blue graph represents the similarity of SNPs compared to Col-0, respectively. Genes underlying this locus are listed in the lower panel and the location of SNPs is indicated with black dashed lines. The bottom graph zooms in the *locus* containing ExAD, and the location and corresponding –log10(P-value) score of the SNPs are shown accordingly using the GWAPP web application (Alonso-Blanco et al., 2016; Seren et al., 2013). The associations above the Bonferroni-corrected threshold are indicated by the red line.

To study the genetic variation in the selected *locus*, we focused on a region that was in linkage disequilibrium with the identified SNPs, spanning approximately 16000 bp containing all the identified SNPs in the candidate *locus*. Subsequently, we conducted a sequence alignment analysis using the Arabidopsis HapMap accessions sourced from the 1001 genome database (Weigel and Mott, 2009) (Figure 2C). Several regions were pinpointed to exhibit missing data or gaps, along with reduced overall similarity, which notably correlated with elevated levels of natural variation as deduced from sequence alignments across all accessions, when compared to Col-0 (Figure 2C, top three panels). Most of the low similarity peaks and significant SNPs with LOD score above the Bonferroni-corrected threshold in Chr3 were found in the *Extensin arabinose deficient transferase* (ExAD, *AT3G57630*) genomic region (Figure 2C, lower panels). We analyzed natural variation in the *ExAD* gene by using 3 significant SNPs (at position 21339391, 21340210 and 21342174) among the 345 Arabidopsis HapMap accessions (Figure S5A and Supplementary Dataset1), and two major haplotypes (Haplotype1 and Haplotype2) for the SNPs within ExAD gene were classified. Accessions with Haplotype1 display significantly higher RVA values after 23 h salt treatment in SITA than Haplotype2 accessions (Figure S5B and Supplementary dataset1). We selected 3 accessions from each haplotype to test whether the expression of *ExAD* changed under salt treatment. Overall, we observed the expression of *ExAD* in the accessions already varied under control conditions (Figure S5C). However, 48 h salt treatment did not cause changes in *ExAD* expression related to specific haplotype and two out of the three SNPs for the ExAD gene are located at exons (Exon 4 and 11) (Figure S5A and S5C), implying that the effect of the SNPs might not affect transcription in response to salt stress.

### Extensin Hyp-Araf_1-4_ arabinosyltransferase activity is required for SITA and halotropism

To further characterize the function of ExAD in regulating root directionality in salt we assessed both the RVA and the root growth rates in SITA of two Arabidopsis T-DNA knockout mutant lines [*exad1-1* (SAIL_843_G12) and *exad1-3* (SALK_204414C) (Figure S6A) that have been previously characterized (Møller et al., 2017). Seedlings were transferred to agar plates and gravistimulated with or without 100 mM NaCl, and quantified for root growth and bending after 24, 48 and 72 h. In both *exad1-1* and *exad1-3* mutants, we observed that the roots followed the direction of the gravity vector more closely in response to salt stress, showing less pronounced inhibition of root bending (Figure 3A) during salt treatment compared to Col-0 (lower RVA values). No difference in root direction was found under control condition when compared with Col-0 seedlings (Figure 3B). Consistently, when Col-0 and *exad1* mutant seedlings were exposed to a salt gradient, the *exad1* mutants exhibited a significantly reduced halotropic (salt avoidance) response (Figure 3C, S6B). Moreover, adult *exad1-1* mutant shoots displayed enhanced Na and K accumulation in salt compared to the wildtype in a hydroponic system (Figure 3D, S6C and S6D).

**Figure 3.**
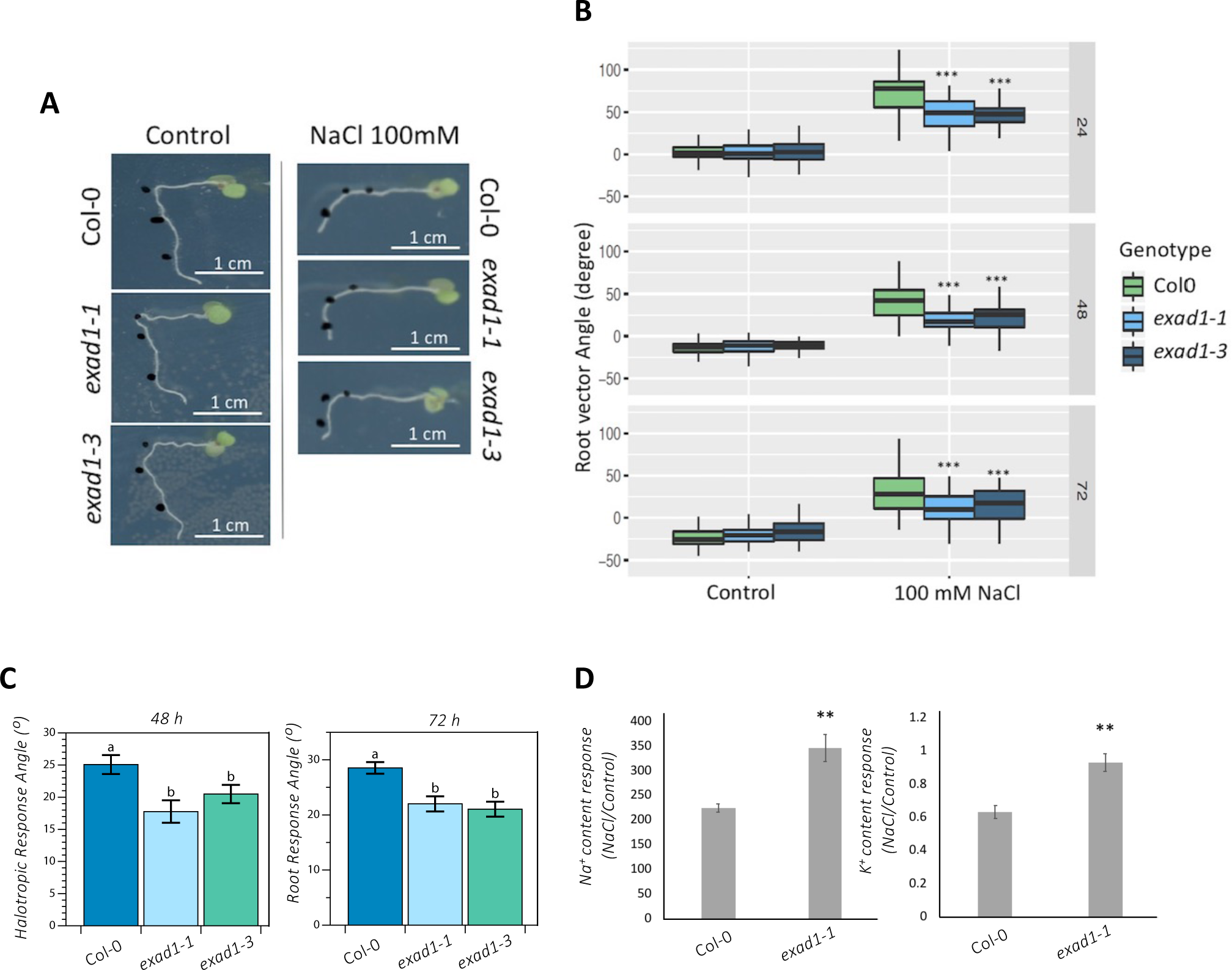
Extensin Hyp-Araf_1-4_ arabinosyltransferase ExAD are involved in modifying root vector angle in salt stress. **A,** Representative image of Col-0, *exad1-1* and *exad1-3* in SITA. The black dots indicate the location of root tips at 0, 24 and 48 h after transferring seedlings to the treatments (with or without 100 mM NaCl). **B**, Quantification of root vector angle of Col-0, *exad 1-1* and *exad1-3* mutants in SITA. Four-day-old seedlings were transferred to plates with or without 100 mM NaCl for 24, 48 and 72 h. Values represent means ± SE from 50 seedlings from 10 plates, each plate containing 5 seedlings. Data in (B) are representative of two independent experiments. Values represent means ± SE from 7 biological replicates (plates) and 10 technical replicates (seedlings). Error bars represent SE of the mean. Statistical analysis was done using two-way ANOVA with contrasts post-hoc. Asterisks indicate statistically significant differences compared to Col0 (p-values ***P< 0.001; **P < 0.01; *P < 0.05). **C**, Quantification of relative root angles on root halotropic response, analyzed in 5-day old seedlings treated with/without NaCl after 48 and 72 h on a gradient, show a reduced response of *exad1* mutant lines. Statistical analysis (n=48) was performed with Shapiro-Wilk test (p < 0.05) followed by non-parametric Kruskal-Wallis test to determine group differences. Letters denote statistically significant differences according to Dunn’s test (p < 0.05). **D**, Na^+^ content response (NaCl treatment/Control treatment) and K^+^ content response were measured in shoots of Col-0 and *exad1-1* mutant plants. Three-week-old plants were grown in hydroponic systems and were transferred to NaCl (150 mM) or control (0 mM) solutions for 4 days. The shoots were then harvested for ion measurement, and the data were normalized by fresh weight. Values represent means ± SE of 4 biological replicates, each containing at least 2 shoots from different plants. Statistical analysis was performed using Student’s T-test. Asterisks indicate statistically significant differences compared to Col-0 (**P < 0.01).

Because similar growth rates were observed across the lines at the different time points analyzed (Figure S6E, S6F), it is likely that the root bending differences detected in salt treated seedlings are specifically linked to the functionality of the ExAD, rather than being influenced by alterations in growth patterns. To test this hypothesis, we examined mutants deficient in XYLOGLUCAN ENDOGLUCANASE 113 (XEG113), an enzyme known to catalyze the addition of the third arabinose residue (Hyp-Araf_3_), which precedes the action of ExAD on extensins (Gille et al., 2009) (Figure S6G, Table S4). Both *xeg113-1* and *xeg113-2* mutants showed a slightly less pronounced root bending (lower RVA values) compared to the corresponding control, but the effect was only visible at 24 h of salt treatment (Figure S6H) and is accompanied with a slight change in root growth rate (Figure S6I). While β-Arabinosyltransferases such as HPAT, RRA, and XEG113 exhibit a broad range of substrates (Matsubayashi, 2014), ExAD stands out as the sole arabinosyl transferase responsible for mediating the addition of α-Araf to Hyps. To date, ExAD is considered specific to the regulation of extensins (Møller et al., 2017; Petersen et al., 2021). This specificity could potentially explain the observed difference between XEG113 and ExAD in controlling root bending responses under salt stress, being the latter more specific for the extensin modification mediated salt dependent responses.

### Salt increases ExAD-dependent arabinosylation signal in Arabidopsis seedlings

To further explore the effects of salt treatment on the extensin repeat side-chains (Hyp-Araf_1-4_) and to detect the overall changes of cell wall polysaccharides, a qualitative survey of cell wall modifications was performed by using the <u>Co</u>mprehensive <u>M</u>icroarray <u>P</u>olymer <u>P</u>rofiling (CoMPP) analysis as described in (Moller et al., 2007). Col-0, *exad 1-1* and *exad 1-3* seedlings were treated with salt (100 mM NaCl) or control (0 mM NaCl) for 48 h before harvesting samples for the alcohol insoluble residues (AIR). CDTA or NaOH were used to perform sequential extractions to obtain the CDTA (pectin-enriched) or the NaOH (hemicellulose-enriched) fractions (Fangel et al., 2021). We quantified signal intensities derived from the recognition of cell wall epitopes through a selection of monoclonal antibodies (mAbs) (Table S5). Overall, we found that in Col-0 and in both *exad 1-1* and *exad 1-3* seedlings salt affects the signals of antibodies involved in the recognition of pectin moieties, extensins and arabinogalactans (Figure S7, Table S6). A previous study suggested that LM1, JIM11 and JIM20 antibodies might be specific to the third Hyp-Araf_3_ or higher order arabinosylation of extensin repeat side-chains (Castilleux et al., 2020). However, to date, the exact targets of JIM11 are not known. Here we report that JIM11 yielded no signal in the *exad 1-1* and *exad 1-3* mutants in either salt or control conditions, likely suggesting that JIM11 might be involved in recognizing the ExAD-dependent Hyp-Araf_4_ (Figure 4A, Figure S7, Table S6). To make sure that JIM11 recognition signals were caused by an actual difference in extensin Hyp-Araf_4_ arabynosilation independently on the way AIR samples were prepared we extracted total proteins by using an adapted method from(Leszczuk et al., 2020; Xu et al., 2011). 25 ug of total protein extracts from Col-0, *exad 1-1 and exad1-3* seedlings treated for 48 h with salt (100 mM NaCl) or control were spotted on nitrocellulose membranes and probed with JIM11 (Figure 4B). Interestingly, again no JIM11 signal was detected in the *exad1* mutant alleles. Similarly, no signal was detected in the *xeg113 1-1* and *xeg113 1-2* mutant seedlings (Figure S8A) likely suggesting that both the ExAD– and the XEG-mediated arabinosylations, altering the overall Hyp-Araf_4_ content, are targets of JIM11 recognition. Moreover, we observed a significant increase in JIM11 intensity in salt-treated Col-0 seedlings compared to control conditions (Figure 4B and S8A), suggesting that salt treatment induces an increase in the JIM11 targets. To evaluate the arabinose levels in the cell wall matrix quantitatively, we analyzed the monosaccharide composition of neutral sugars in AIRs extracted from seedlings of Col-0, *exad 1-1*, *exad 1-3* mutants treated for 48 h with NaCl or mock (Figure 4C). Our data show a clear arabinose phenotype in both the *exad 1-1* and *exad 1-3* mutants characterized by a mild reduction (likely being associated to the Hyp-Araf_4_ deficiency). We also detected a more pronounced arabinose reduction in the *xeg113-1* and *xeg113-2* mutant lines likely being dependent on an impairment in addition of both Hyp-Araf_3_ and Hyp-Araf_4_ (Figure S8B). In addition while we could confirm that salt triggers an increase in galactose content and a reduction in xylose (Figure 4C, S8B), a phenotype which seems consistent in all the analyzed lines, we could not detect a significant salt-triggered cellulose reduction (Figure S8C) which has been proposed to be mechanistically linked to changes in galactose content (Yan et al., 2021). Cellulose levels appear to be comparable in the mock treated *exad 1-1*, *exad 1-3, xeg113-1, xeg113-2* and Col-0 seedlings, hence do not seem to correlate with a change in JIM11 dependent signal and/or root bending phenotypes in controls (Figure S8C).

**Figure 4.**
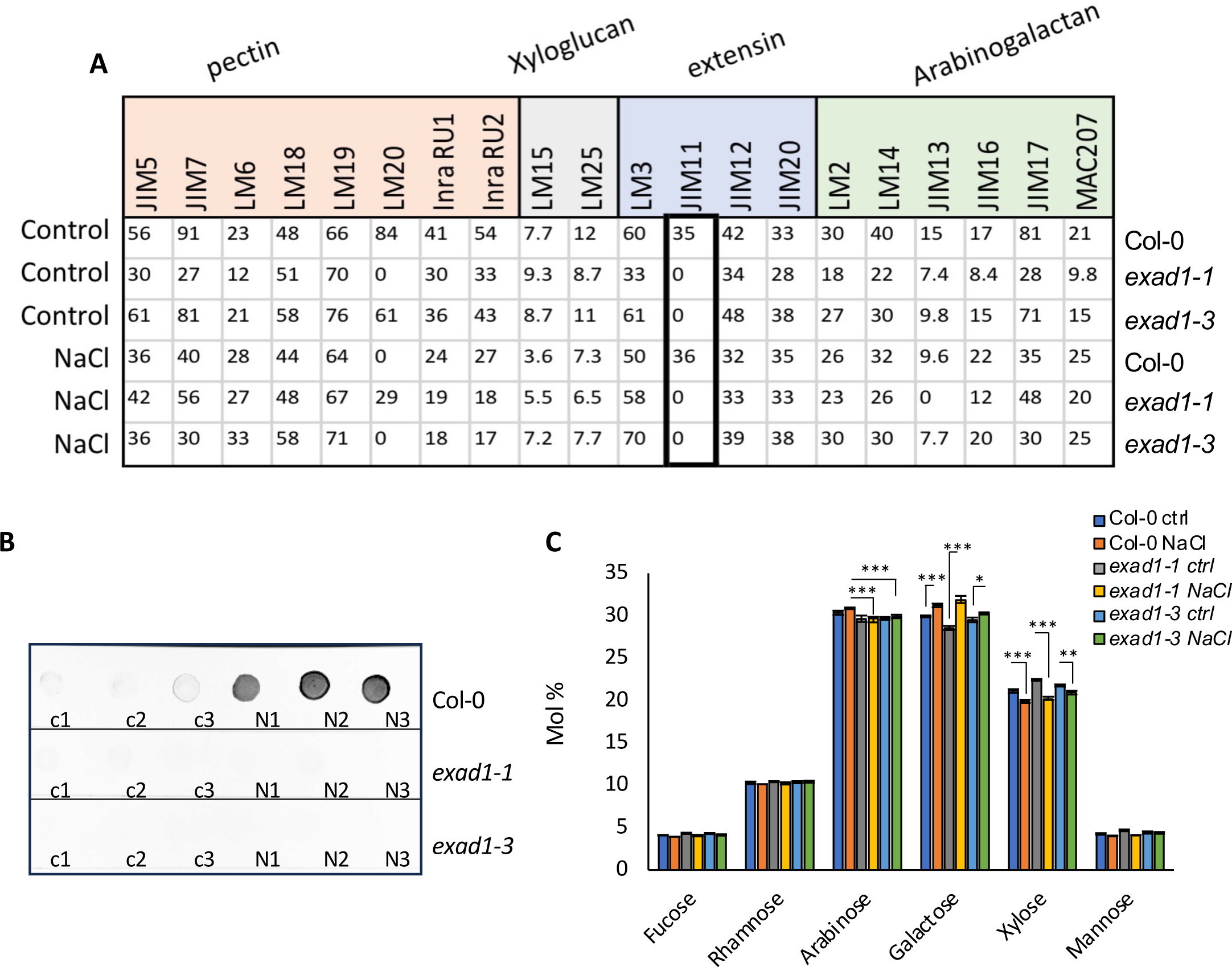
Anti-extensin monoclonal antibody (mAB) JIM11 recognizes the ExAD-mediated arabinosylation. **A,** Comprehensive Microarray Polymer Profiling (CoMPP) analysis quantified mean spot intensities values (Y-axis) showing the cell-wall glycans recognized by a selection of antibodies and carbohydrate binding modules (CBMs) within the pectin-enriched fraction (CDTA extraction). Five-day-old Arabidopsis seedlings of Col-0, *exad1-1* and *exad1-3* mutants were transferred to the liquid treatment medium containing control (0 mM NaCl) or salt (100 mM NaCl) for 48 h then harvested for cell wall alcohol-insoluble residue (AIR) extractions. The list of the antibodies used can be found in Table S5. Values represent means ± SE from 3 biological replicates that were isolated. Full dataset of CoMPP heatmap for all extractions (CDTA and NaOH) are shown in Supplemental Figure S8. **B,** Five-day old seedlings of Col-0, *exad1-1, exad1-3,* were treated with or without 100 mM NaCl for 48 h before harvesting. Total protein were extracted and 25 ug of total protein spotted on nitrocellulose membrane. JIM11 antibody was used to detect the specific Hyp-Ara_4_ signal. Dot-blot im a g e s ar e representative of 2 independent experiments performed each containing 3 biological replicates per treatment [Control (c1, c2, c3), or NaCl (N1, N2, N3)] per genotype. **C,** Five-day old seedlings of Col-0 *exad1-1, exad1-3* mutants were treated with 100 mM NaCl or without salt (Ctrl) for 48 h. Neutral cell wall matrix components were extracted from AIRs derived from 3 biological replicates per treatment/genotype and expressed as Molar Percentage (Mol%). Error bars represent SD of the mean values analyzed for each biological replicate (n= 3). Asterisks indicate statistically significant differences according to Student’s t-test compared to Col-0 NaCl (for Arabinose) or compared to the corresponding controls (for Galactose and Xylose) (*, P< 0.05; **P< 0.01 ***P< 0.001).

### ExAD is required for maintaining root cell wall thickness under salt treatment

To investigate whether the ExAD dependent Hyp-arabinosylation induced by salt would change cell wall structure, we quantified cell wall thickness in root epidermal cells by transmission electron microscopy (TEM). In our analyses we also included xeg*113-1, xeg113-2* mutants in addition to *exad 1-1*, *exad 1-3* and Col-0 control seedlings. Since *ExAD* is highly expressed in the maturation zone (Møller et al., 2017), we examined cell wall thickness in both the elongation and in the maturation zone. When seedlings were treated with NaCl a significant change in cell wall width was detected in both Col-0 and in the *exad1* loss of function mutant alleles (Figure S9, 5A, 5B). Interestingly, the increase in cell wall thickness in response to salt was greater in the *exad1-1 and exad1-3* mutants compared to wild type seedlings, in both the elongation and maturation zones of the root (Figure 5), while the *xeg113* mutant alleles did not show any significantly statistical difference in cell wall thickness compared to Col-0 seedlings (Figure 5B). Also no significant differences between *exad1-1* and Col-0 seedlings were detected in control conditions (Figure S9). Our results suggest that in response to salt stress, cell wall width changes in wild type seedlings and that this event seems to be controlled by ExAD function in a salt dependent manner. To delve deeper into the structural changes of cell wall mechanics and properties in response to salt stress, we utilized a previously described fluorescent mechano-probe (Michels et al., 2022). The mechano-probe’s lifetime, which serves as an indicator of changes in cell wall density and strength, was measured using Fluorescence Lifetime Imaging Microscopy (FLIM). We collected emission spectra of the probe in elongation zone epidermal cells of Col-0, *exad1-1* and *exad1-3* seedlings that had been treated with 100 mM NaCl for 48 hours (Figure 5C). Our analysis revealed that in the absence of *ExAD*, the mechano-probe exhibited a significantly lower lifetime under salt conditions. This observation likely indicates higher cell wall porosity associated with less dense cell walls and reduced cell wall strength. This finding aligns with the TEM analysis, suggesting that in the absence of ExaD, the extensin-dependent crosslinks, which appear to be altered in the presence of salt, affect the properties of the cell walls, resulting in less rigidity and swollen cell walls.

**Figure 5.**
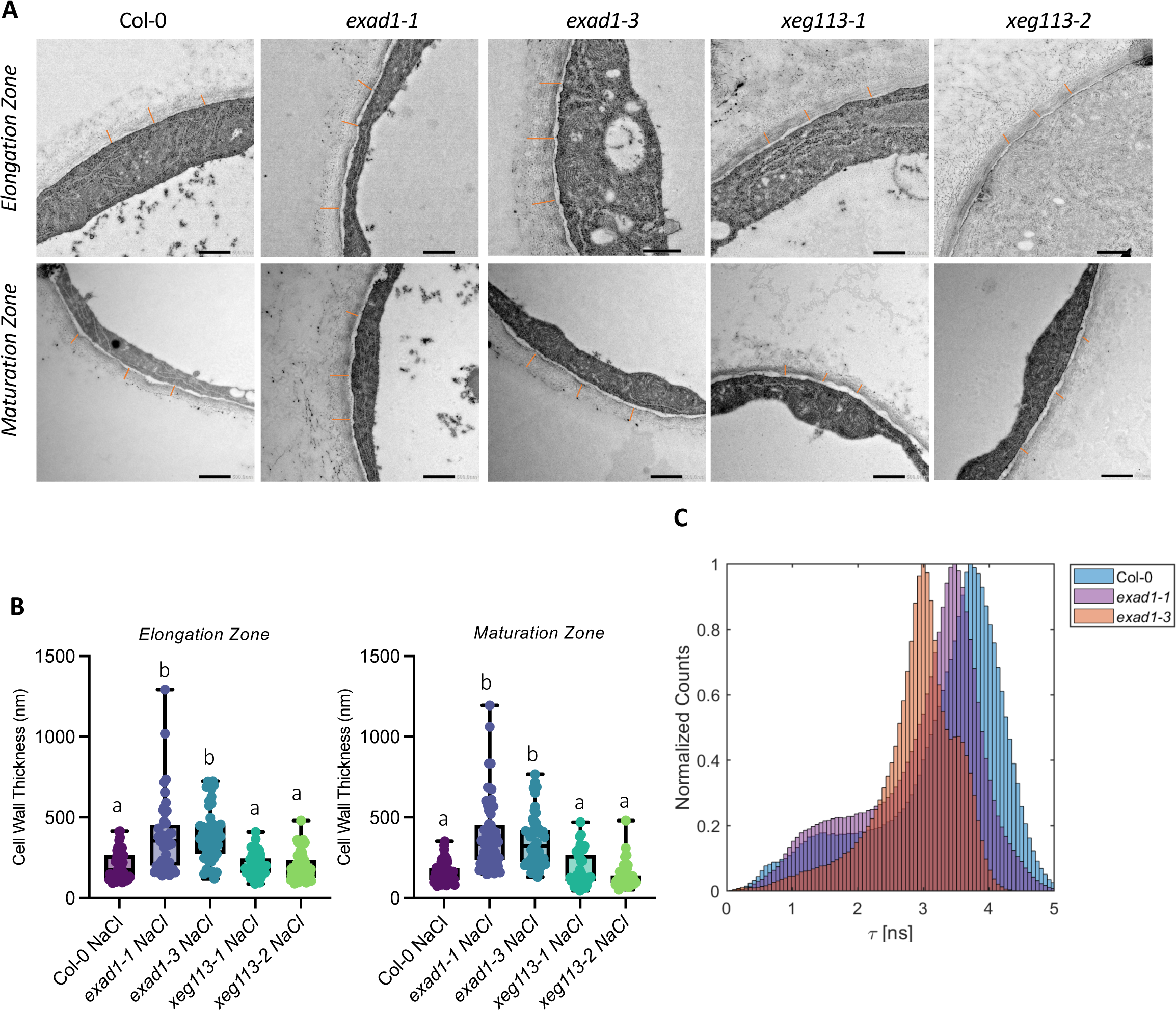
ExAD regulates root cell wall thickness under salt treatment. **A**, Cell wall structure of cells in epidermal layers in the root elongation zone (upper panel) and in the root maturation zone lower panel. Using the SITA system, four-day-old Arabidopsis seedlings of Col-0, *exad1-1*, *exad1-3*, *xeg113-1* and *xeg113-2* mutants were transferred 100 mM NaCl for 48 h before sampling for TEM analysis. Representative images are shown. **B,** Cell wall thickness of epidermal cells or maturation zone cells was measured in three biological replicates, each consisting of 16-20 independent cells. Cell walls were measured at 3 randomized positions on each of the analyzed cells per biological replicate. Letters indicate statistically significant differences between genotypes according to one-way ANOVA and Tukey’s HSD test (α = 0.05). **C**, Effects of salt treatment changes cell wall mechano-chemical properties in an ExAD dependent manner. Col-0 *exad1-1* and *exad1-3* mutant seedlings were treated with 100 mM NaCl for 48 h. The intensity of the mechano-sensing probe described in (Michels et al., 2022) displays altered cell wall porosity in the *exad1* mutant lines detected through FLIM analysis. The seedlings were stained with CWP-BDP and fluorescence emission of elongation zone epidermal cells was captured on a Leica TCS SP8 inverted confocal microscope coupled to a Becker&Hickl TCSPC lifetime module (SPC830). Samples were excited with a 514-nm pulsed laser source (pulse duration < 1 ps) with a repetition rate of 40 MHz and plotted as an average of 3 independent experiments each containing 5 seedlings per genotype (n=15).

## Discussion

Salt stress negatively affects growth and development of roots in most plant species (Julkowska et al., 2014; van Zelm et al., 2020). Roots can change their growth direction dynamically to rapidly avoid a saline environment, a response that can be used as a phenotypic output to study salt signalling pathways (Deolu-Ajayi et al., 2019). In response to salt gradients, roots bend away from high salt, a process called halotropism. The mechanisms of how root halotropic responses are accomplished by redistribution of auxin have been studied in recent years (Galvan-Ampudia et al., 2013; Korver et al., 2020). On the other hand, in uniform salt conditions, little is known about the regulatory pathways and genetic components that are required for salt-induced directional growth (Sun et al., 2008). In this study, we describe a salt-induced tilting assay (SITA) to investigate root directional growth upon gravitropic challenge (gravi-stimulation) in response to salt. We found the SITA response could be modulated by salt, specifically by Na^+^ ions, but not by an equivalent level of osmotic stress (Figure 1B, D, S1A). This finding is akin to the halotropism response, which is also specific to Na^+^ ions as demonstrated by previous studies (Galvan-Ampudia et al., 2013; Deolu-Ajayi et al., 2019). We identified genetic components that may control root directional growth in SITA by using GWAS, by screening a collection of 345 natural Arabidopsis accessions. Both phenotype data and associations presented here (all raw phenotyping data available on request and to be deposited in a public repository upon publication) are a useful resource for follow-up approaches to understand Na^+^-specific responses in roots. Here, we focused on the characterization of one of these identified genes, encoding for ExAD, an arabinosyl transferase involved in adding the fourth arabinofuranose (Hyp-Araf_4_) on the continuous stretches of hydroxyprolines present on cell wall localized glycoproteins, such as the plant extensins. Extensins are repetitive hydroxyproline-rich O-glycoproteins (HRGPs) that have been suggested to modulate plant primary cell wall architecture (Hijazi et al., 2014). In our study, the *exad1-1* and *exad1-3* loss of function mutants showed reduced salt-induced modulation of root gravitropic bending and halotropism response (Figure 3), reduced levels of Na and K in roots, and a significantly thicker cell wall structure under salt conditions (Figure 5B, C), which may be linked to possible extensin cross-linking defects in the *exad1* mutants. Well-cross-linked extensins are known to play crucial roles in supporting and maintaining cell wall architecture and cell wall formation, regulating plant growth and development, and to contribute to disease and wounding resistance (Lamport et al., 2011; Cannon et al., 2008; Lamport and Várnai, 2013). Various types of cross-links, that are built with different cell wall polymers, can be classified into homopolymeric and heteropolymeric cross-links (Mishler-Elmore et al., 2021). The heteropolymeric cross-links have been found between cell wall polymers, including pectin and cellulose (Pérez García et al., 2011), pectin and extensin, pectin and hemicellulose, AGP and extensin, and extensin and lignin, as summarized in (Mishler-Elmore et al., 2021). Extensins can also form homopolymeric cross-links, which require correct Hyp-arabinosylation. The extensin peroxidases PEROXIDASE9 (PRX9) and PEROXIDASE40 (PRX40) were proposed to contribute to extensin cross-linking during pollen development in Arabidopsis (Jacobowitz et al., 2019). Interestingly, also arabinosylation, in particular the ExAD-dependent Hyp-α-arabinosylation to form Hyp-Araf_4_, may be crucial for extensin crosslinking *in vitro* (Chen et al., 2015). A biotic stress study suggested that extensin arabinosylation may be involved in cell wall formation and maintenance, and to protect against infection by the oomycete *P. parasitica* (Castilleux et al., 2020). Most recently, it has been hypothesized that defects in extensin arabinosylation in the Hyp-Araf mutants cause deficiency in extensin cross-linking, which may lead to cell wall architecture changes and eventually decrease plant defenses (Castilleux et al., 2021; Tan and Mort, 2020). Here, using TEM analysis we show the involvement of ExAD-dependent Hyp-α-arabinosylation in regulating root cell wall structure under salt treatment (Figure 5B), which seems to be linked to alterations of root bending in response to salinity stress and the ability of roots to avoid high salinity by change in growth direction (halotropism) (Figure 3). *exad* mutant roots of plants exposed to salinity had higher levels of Na and K, suggesting uptake of ions in general could be increased in the mutants. Moreover by using a cell wall mechano-probe (Michels et al., 2022) we display that the cell wall density and porosity depends on the function of ExAD and is thus likely linked to the salt triggered ExAD-mediated Hyp-α-arabinosylation. Salt stress induces several cell wall-localized responses, such as changes in polysaccharide deposition, changes in pectin properties and microfibril orientation, which could compromise cell wall function (Byrt et al., 2018). Reduction of cellulose content has been reported before (Endler et al., 2015; Yan et al., 2021), but under our experimental conditions we could not detect cellulose changes. It is possible that at the analyzed time point (48h) upon salt stress, cellulose content reduction is not yet evident while the extensin-mediated phenotype can already be quantified. Recently, the cell wall has indeed been hypothesized to be crucial for salt tolerance and salt signalling pathways (Feng et al., 2018; Gigli-Bisceglia et al., 2022). The *Catharanthus roseus* receptor-like kinase 1-like (CrRLK1L) protein kinase subfamily member FERONIA (FER) and the HERK1/THE1 combination have been suggested to be responsible for triggering several salt response signalling pathways, which depend on the detection of salt-induced pectin modifications (Gigli-Bisceglia et al., 2022; Feng et al., 2018). As a chimeric class of extensins, Leucine-rich repeat extensins (LRXs) members LRX1 and LRX2 were suggest to be involved in root hair formation and cell morphogenesis via mediating cell wall development (Baumberger et al., 2003, 2001). LRX3/4/5 were shown to be required for plant salt tolerance, via binding of Rapid Alkalinization Factor (RALF) peptides and their direct interaction with FER to transduce the cell wall signals in response to salt stress (Zhao et al., 2018). The *exad* loss of function mutants were reported to have shorter root hairs (Møller et al., 2017), similar to other extensin Hyp-β-arabinotransferase mutants, such as the *rra* (responsible for Hyp-Araf_2_) and *xeg113* (responsible for Hyp-Araf_3_ formation) (Egelund et al., 2007; Velasquez et al., 2011; Gille et al., 2009), while *hpat* mutants (responsible for Hyp-Araf_1_) were reported to have defects both in cell wall thickness and root hair elongation (Ogawa-Ohnishi et al., 2013; Velasquez et al., 2015). Whether the salt triggered ExAD dependent phenotypes are linked to reduced root surface area due to shorter hairs, previously suggested to alter salt sensitivity (Robin et al., 2016), is yet unclear. Here we show that changes in root bending response in salt stress seems to be likely dependent on changes in cell wall width and resulting mechanical changes. We now suggest that extensin arabinosylation constitutes another type of cell wall modification, which is not necessary for root growth but crucial for the directional response to salt stress. Extensins, as structural cell wall glycoproteins, appear to play a vital role in maintaining the normal physical structure of the cell wall under stress conditions. It can be hypothesized that salt-induced cell wall modifications and the associated changes in turgor pressure activate the ExAD enzyme, which is involved in increasing extensin arabinosylation to enhance cell wall strength under challenging conditions. In contrast, in physiological conditions when the cell wall is unchallenged, enhanced crosslinking is not required, making ExAD dispensable. However, in situations where cell walls are under stress, the function of ExAD becomes essential to counteract changes in the structural integrity of the cell wall. Hence, in its absence, the cell wall might become weaker, more hydrated, and thicker (Figure 5). We propose that salt-induced extensin arabinosylation is a critical mechanism for reinforcing the cell wall and mitigating the damage caused by Na^+^ ions, enabling roots to respond effectively to salinity stress. In summary, we found that salt stress affects cell wall composition and structure in an ExAD-dependent manner, revealing that ExAD α-arabinosylation of extensin to yield Hyp-Araf_4_ is required to build up extensin cross-linking and to modify cell wall structure during salt stress. As such, our SITA time lapse assay of root growth of a set of natural accessions in salt has revealed that cell wall modification process(es)/changes are a crucial aspect of root responses to salinity, similar to their importance for biotic stress perception (Gigli-Bisceglia and Testerink, 2021). How this process connects to other reported salt-induced cell wall modifications and how they together function in root responses and plant performance in saline conditions, remains to be established.

## Material and Methods

### Plant materials and growth conditions

All the seeds used in this study (listed in Table S1 and Table S4) were propagated under long day (16 h light/ 8 h dark cycle), 21°C, 70% humidity. *Arabidopsis thaliana* seedlings of all the phenotypic assays in this study were grown under long day conditions (16 h under white LED light/ 8hrs dark cycle), 21°C, 70% humidity. Seeds were sterilized by 30% bleach solution and stratified at 4°C for 3 days in dark before they were sown on agar plates. Seeds of mutant plants *exad1-1* (SAIL_843_G12), *exad1-3* (SALK_204414C) were previously characterized (Møller et al., 2017). *xeg113-1* (SALK_151754) and *xeg113-2* (SALK_066991) were ordered from NASC. Detailed information of seeds is listed in the Table S4. For SITA, ½MS medium containing 0.1% 2(N-morpholino) ethanesulphonic acid (MES) buffer (Duchefa) and 0.5% Sucrose (Duchefa), pH 5.8 adjusted with KOH, 1% Daishin agar (Duchefa) was used. For salt treatment, NaCl was added after adjusting pH to 5.8. In the salt specificity experiment, treatments were Control, 100 mM NaCl, 100 mM NaNO_3_ 200 mM Sorbitol, 100 mM KCl. In the salt dose response experiment, treatments were Control, 75, 100, 125 mM NaCl as indicated in the figure legend. In all other experiments, treatments were Control and 100 mM NaCl. For gene expression analysis of ExAD1 in T-DNA mutants (*exad1-1* and *exad1-3*), seeds were germinated and grown for eight days before harvesting, on agar plates containing ½ MS medium with 0.5% sucrose, 0.1% MES (Duchefa), 1% Daishin agar (Duchefa), pH 5.8 adjusted with KOH. For gene expression analysis of ExAD in tested accessions from two haplotypes, seeds were germinated and grown in a flask containing ½ MS medium with 0.5% sucrose, shaking on shaker with the speed of 130 rpm/min under long-day condition. Four-day old seedlings after germination were transferred to control or NaCl (100 mM) for 48 h before harvesting. For dot blot assays, seeds were germinated and grown in a flask containing ½ MS medium with 0.5% sucrose and 0.1% M.E.S. shaking on shaker with the speed of 130 rpm/min under long-day condition. Four-day old seedlings after germination were transferred to control or NaCl (100 mM) for 48 h before harvesting. For CoMPP analysis, seeds were germinated and grown in a flask containing ½ MS medium with 0.5 %sucrose and 0.1 % MES. shaking on shaker with the speed of 130rpm/min under long-day condition. Four-day old seedlings after germination were transferred to control or NaCl (100 mM) for 48 h before harvesting. For the ion content analysis in hydroponic system (https://www.araponics.com/): 3-week-old Arabidopsis plants of Col-0 and *exad1-1* mutant were grown in hydroponics and treated with NaCl (150 mM) or Control (0 mM) for 4 days. Liquid medium was used (½ MS including vitamins, 0.1% MES, pH 5.8). Shoots were harvested and approximately 150 mg of fresh plant material was harvested and rinsed with MQ water (3 times). Next, the shoots were submerged in a solution of 10 mM CaCl2 for 5 mins and washed with MQ water; submerged again with a solution of 10 mM EDTA for 5 mins and washed with MQ water (in total 3 times). All samples were dried at 100° C for 48 h prior to ion measurement. Ion measurements were performed by ICP-MS as described in (Danku et al., 2013) at the Ion omics Facility, University of Nottingham, UK.

### SITA

*Arabidopsis thaliana* seeds were dry sterilized with 40 ml thin bleach and 1.2 ml 37% HCl in a desiccator for 3 h, followed by at least 3 h in laminar flow to remove the extra surface bleach. Then seeds were stratified in 0.1% Daishin agar solution at 4°C in the dark for 3 days before sowing. After stratification, seeds were germinated on fresh half strength (½) MS medium 1% agar plates that were placed in 70-degree angle racks for 4 days (80 seeds per plate). Four-day-old Arabidopsis Col-0 seedlings were transferred to ½ MS agar plates containing different treatments. After transferring seedlings to treatments, all plates were rotated 90 degrees clockwise simultaneously to apply a gravistimulus and placed in vertical (90 degree) racks. In the time-lapse system, all plates were captured every 20 minutes by infrared photography for at least 24 h. Alternatively, root tips were marked up at 24 h, 48 h and 72 h as indicated in figure legends.

### Halotropic response analysis

Halotropism assays on salt gradients were performed as described before (Galvan-Ampudia et al., 2013; Deolu-Ajayi et al., 2019). *Arabidopsis thaliana* seeds (Col-0, *exad1-1* and *exad1-3*) were dry sterilized and stratified in 0.1% Daishin agar solution at 4°C in the dark for 3 days before sowing. After stratification, seeds were germinated on fresh half strength (½) MS medium 1% agar plates with 0.5% sucrose in 90-degree angle racks for 5 days. After 5 days, a 45° cut-out of the medium was made at 0.2 cm below the root tips and replaced with medium supplemented with/without 200 mM NaCl. Plates were scanned with an Epson Perfection V800 Scanner at 48 and 72 hours after the treatment. The images were analyzed with Smartroot.

### Root phenotypic quantification and data processing

In the time-lapse system, images of plates were captured every 20 min in jpg format, at 690dpi. Alternatively, images of plates were scanned with an Epson Perfection v800 Photo scanner at 400dpi in jpg format every day. All images were improved to black/white in tiff format, and each root traced and quantified in ImageJ, SmartRoot plugin. Two root traits were used for quantifications: Root vector angle (RVA) is measured from the beginning of the root to root tip as shown by the white arrow. Root tip direction (RTD) is indicated by the last 10% of the root tracing length (Figure 1A). The temporal trait K^N/C^ (response in fitted rate of exponential decay) was obtained by fitting a model of exponential decay (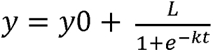) to the changes in root tip direction (y) over time (t) for every accession in both salt and control conditions. In this model, y□ represents the lower asymptote, L is the initial decrease in angle from the starting value to the lower asymptote, and k is the rate of exponential decay. The parameter k was used in the GWAS as a response variable between salt and control conditions. The time-lapse system has the capacity of maximum eight plates vertically in a row and the camera (Canon 1200D) on the rail in front of plates that can move and take pictures. For each plate, the images were firstly traced manually with SmartRoot for the last time-point image, and then a SmartRoot plugin –automatic tracing-was used to trace images back at earlier time points. All the nodes during tracing were copied from the previous images to the continuous next image, and simultaneously detect the pixel intensity to remove the extra nodes from root tip. After automatic tracing, nodes in each image were inspected and adjusted manually if needed. SmartRoot was used in both time-lapse and scanning system tracing and output data into csv file, available at https://smartroot.github.io/ (Lobet et al., 2011). R was used for data and graphs processing (R scripts are available on request).

### Genome Wide Association Study (GWAS)

Root SITA response traits of the HapMap accessions were collected using a time-lapse system. In the phenotyping experiment for GWAS, 5 replicates (seedlings) on each plate were used to calculate the average used as input. The time-lapse system has the capacity of maximum eight plates vertically in a row, and Col-0 was repeated in each round of timelapse scanning as the standard control. The phenotypic data was used in concert with SNP markers from the 1001 genome project, and Genome Wide Association Study (GWAS) was performed by two association mapping tools: GWAPP (Alonso-Blanco et al., 2016; Seren et al., 2012) and R using the EMMAX and the ASReml R package (version 3) (Butler et al.). The GWAS script included terms for correcting population structure and the kinship matrix (Korte et al., 2012). To determine population structure, PCA was performed using the factoextra R package (Kassambara and Mundt, 2020). Although the total number of SNPs encompassed 4,285,827 SNPs, only SNPs with minor allele frequency > 0.05 were considered for validation. Therefore, the Bonferroni threshold was determined by the – log_10_(p-value / # SNPs) for SNPs with minor allele frequency > 0.05. This corresponded to – log_10_ (0.05/1,753,576) = 7.55. The broadsense heritability was calculated using MVApp (Julkowska et al., 2019). The GWAPP web application is available at http://gwapp.gmi.oeaw.ac.at/ (Table S2). The data followed a normal distribution and the threshold for significant SNPs was determined by correcting p < 0.1 with Bonferroni correction. Total number of SNPs encompassed 204741 SNPs. Haplotype analysis was performed for the ExAD gene using 3 significant SNPs found in GWAS. The genotypes of the Hapmap accessions were sorted according to the allele of each SNP, which revealed 2 haplotype groups.

### Sequence comparisons among accessions

Sequence information of HapMap accessions were obtained from the 1001 Genome Project (Weigel and Mott, 2009), available at http://signal.salk.edu/atg1001/3.0/gebrowser.php. Sequences were aligned with ClustalO, and the comparisons are presented in three plots including missing data, gaps and similarity by Gnu-plot software package(Julkowska et al., 2016).

### T-DNA insertion line genotyping

Leaf material of 2-week-old plants was collected for DNA isolation. Tissue was ground in liquid N2, incubated with STE/Lysis buffer (100mM Tris pH 7.5, 2% SDS, 10 mM EDTA) at 65 °C for at least 20 minutes, precipitated with Ammonium Acetate (NH4Ac). Supernatant was collected and incubated with 2-propanol and spun at a max speed. The DNA pellet was resuspended in water for PCR. Primers and SALK numbers are listed in Table S4.

### Gene expression analysis

For gene expression analysis of ExAD in T-DNA mutants (*exad1-1* and *exad1-3*), seeds were germinated and grown on agar plates. Eight days after germination, seedlings were harvested, frozen by using liquid N_2_, and ground into a fine powder with a paint shaker for further RNA extractions. Total RNA was extracted and purified in TRIPure (Sigma), followed by chloroform and isopropanol cleaning steps. RNA pellet was washed by 70% ethanol and followed by DNase treatment (Ambion). cDNA synthesis was performed using iScript cDNA synthesis kit (Bio-Rad). Reverse transcription qPCR (RT-qPCR) was carried out using Bio-Rad CFX96 system. Relative expression was calculated using the reference gene AT2G43770 and At2g43770 individually, and the average was used. Values represent means ± SE from 9 tubes from 3 plates, each tube containing 2 seedlings. For gene expression analysis of ExAD in tested accessions from two haplotypes, seeds were germinated and grown in a flask, shaking on shaker with the speed of 130rpm/min under long-day condition. Four days after germination, seedlings were transferred to control or NaCl (100 mM) for 48 hrs before harvesting. Seedlings were harvested, frozen by using liquid N_2_, and ground into a fine powder with a paint shaker for further RNA extractions. Five biological replicates for each treatment and each genotype were used. Total RNA was extracted and purified by using NZY Total RNA Isolation kit (NZYTech). cDNA synthesis was performed using iScript cDNA synthesis kit (Bio-Rad). The qRT-PCR was carried out using a CFX Opus 384 Real-Time PCR System (Bio-Rad). Relative expression was calculated using the reference gene *AT2G43770*. Primers used for gene expression analysis were listed in Table S4.

### Sequential extraction and Comprehensive Micro Array Polymer Profiling (CoMPP) analysis

Five-day-old Arabidopsis seedlings of Col-0, *exad1-1* and *exad1-3* mutants were transferred to the treatments containing (0 mM NaCl) as control, or 100 mM NaCl for 48 h before sampling. Seedling samples were harvested, flash frozen, and lyophilized. The dried material was ball milled, and the alcohol insoluble residue (AIR1) was removed by incubating the material first for 30 min in 80% ethanol, and then for 30 min in 70% ethanol at 95 °C, and finally in chloroform: methanol (1:1) for 5 min at room temperature before washing with acetone. This sequence of incubations and washes constitutes the AIR1 treatment mentioned above. Starch (AIR2) was then removed according to (Møller et al., 2017). AIR was extracted sequentially first with 50 mM CDTA (1,2-Cyclohexylenedinitrilotetraacetic Acid) (CDTA; Merck Darmstadt, Germany) followed by 4 M NaOH, 26.5 mM NaBH_4_ (Merck Darmstadt, Germany). The resulting fractions (primarily pectin and hemicelluloses, respectively) was dotted on nitrocellulose using a piezoelectric array printer (Marathon, Arrayjet, Edinburgh, United Kingdom) resulting in microarrays with all the samples extractions. The microarrays were then probed with a selection of antibodies and carbohydrate binding modules (CBMs). A list of the antibodies used can be found in Table S5. The microarrays were blocked with skimmed milk, probed with CBMs or monoclonal antibodies specific to glycan epitopes, and then visualized using the appropriate secondary antibodies as described in (Møller et al., 2017). Spot intensities are quantified from 256-step gray scale scans of the microarrays utilizing array detection software (Array-Pro Analyzer v 6.3, Media Cybernetics, Rockville, Maryland, United States) and data are presented on a relative scale with 100 assigned to the most intense spot. The extracted CoMPP fractions were dialyzed against demineralized water using Visking® 12-14 kDa dialysis tubing (45 mm) prior to determination of the monosaccharide composition.

### JIM11 dot blot assay

Five-day-old seedlings were transferred to treatments of control or NaCl (100 mM) for 48 h before harvesting. Samples were frozen using liquid N_2_, ground into a fine powder with a paint shaker. At least 3 biological replicates for each treatment and each genotype were used. Total proteins were extracted in buffer: 0,5 M Tris-HCl (pH 6.8), 0.037 g/L EDTA (Na_2_ EDTA•2H_2_O), 2% SDS. Samples were boiled at 98 °C for 3 min. After a short centrifuge (15 mins, 13.000 xg), supernatants were collected, and total proteins quantified though Bradford assay. 25 µg of total proteins were spotted on nitrocellulose membrane. Membranes were dried overnight an RT. Blocking (5% BSA in PBS) was performed for 1 h. Membranes were incubated for 1 h with primary antibody JIM11 (Agrisera) (1:500, 5% BSA in PBS). Membranes were incubated with secondary antibodies rat (IgG) (1:6000, 5% BSA i n PBS) and after 3 x washing steps (PBST), chemiluminescence was induced by Clarity Western ECL Substrate (BIO-RAD) and detected with ChemiDoc imaging system (Biorad).

### Transmission electron microscopy (TEM)

Four-day-old seedlings were transferred to treatment agar plates supplemented with 100mM NaCl or 0mM as control for 48 h before sampling in SITA system. Specimen was cut into small pieces of about 1mm^3^ at the selected zones (elongation zone and maturation zone) before fixation. Root pieces were submerged and incubated with (2.5% glutaralehyde + 2% paraformaldehyde in 0.1M phosphate/citrate) buffer for at least 1 hour at room temperature. Specimen was washed at least 6 times for 10 mins in 0.1M phosphate/citrate buffer (washing buffer). 1% osmium tetroxide in 0.1M phosphate/citrate buffer was added and incubated for 1 hour at room temperature. Specimen was washed at 3 times for 10 minutes in MQ. Next, specimen was dehydrated in a graded ethanol series: 30% for 5 mins, 50% for 5 mins, 70% for 5 mins, 80% for 5 mins, 90% for 5 mins, 96% for 5 mins, 100% for 10 mins, and infiltrated with resin (spurr’s resin) 1:2 resin:ethanol for 30 mins, 1:1 resin:ethanol for 30 mins, 2:1 resin:ethanol for 30 mins, 100% resin for 60 mins or overnight at room temperature. Leica EM Rapid was used for trimming (pre-sectioning) of the sample and Leica Ultramicrotome U7 was used for sectioning and attaining the sections. Sections were viewed using a JEOL 1400 TEM operating at 120 kV.

### Cell wall analysis

Five-day-old seedlings of Col-0, *exad1-1*, *exad1-3* and Col-0, *xeg113-1, xeg113-2* were treated with 0 mM or 100 mM NaCl. Seedlings were harvested after 48 h of salt treatment. Arabidopsis seedlings were lyophilized and ball-milled in a Retsch mixer mill. All samples were extracted three times with 70 % ethanol and three times with 1:1 (v:v) chloroform:methanol in the Retsch mill, washed with acetone and dried in a vacuum concentrator. The alcohol insoluble residue (AIR) was weighed out in 2 ml screw cap tubes and used for extraction of neutral cell wall sugars and cellulose as described (Yeats et al., 2016). High-performance anion-exchange chromatography with pulsed amperometric detection (HPAEC-PAD) was performed on a biocompatible Knauer Azura HPLC system, equipped with an Antec Decade Elite SenCell detector. Monosaccharides were separated on a Thermo Fisher Dionex CarboPac PA20 column with a solvent gradient of (A) water, (B) 10mM NaOH and (C) 700mM NaOH at 0.4 ml/min flow rate. 0 to 25 min: 20% B, 25 to 28 min: 20 to 0% B, 0 to 70% C, 28 to 33 min: 70 % C, 33 to 35 min: 70 to 100% C, 35 to 38 min: 100% C, 38 to 42 min: 0 to 20% B, 100 to 0% C, 42 to 60 min: 20% B.

### Synthesis of the Cell Wall mechano-probe

The cell wall mechano-probe for FLIM analysis, as detailed in (Michels et al., 2022), was employed in this study. The chemical synthesis method differed from the previous publication and its procedure is outlined below. Step 1: Synthesis of 4-(azidomethyl)benzaldehyde. 4-(bromomethyl)benzaldehyde (5g, 25 mmol) and NaN_3_ (2.5 g, 38 mmol) were added to 50 mL of DMF in a round bottom flask. The solution was left stirring at 60 LC for 1.5 hours. After cooling the solution was diluted with 250 mL of ethyl acetate and washed with 2×250mL of water. The resulting organic layer was dried with MgSO_4_, filtered, concentrated and dried under vacuum to yield 4-(azidomethyl)benzaldehyde (3.76 g, 93% yield). 1H NMR (400 MHz, CDCl3) δ 10.02 (s, 1H), 7.90 (d, J = 8.1 Hz, 2H), 7.49 (d, J = 8.5 Hz, 2H), 4.45 (s, 2H).

Step 2: Synthesis of N_3_-BODIPY mechano-probe. To a 1L 3-neck round bottom flask 4-(azidomethyl)benzaldehyde (2.0 g, 12.41mmol) was added followed by 500 mL of anhydrous dichloromethane and 2-methylpyrrole (2.092 mL, 24.82 mmol). After sparging the solution with N_2_ for 30 minutes trifluoroacetic acid (500 µL, 6.2 mmol) was added. The reaction mixture was left stirring for 2 hours followed by the addition 2,3-Dichloro-5,6-dicyano-1,4-benzoquinone (2.82 g, 12.41 mmol) after which the mixture was sparged with N_2_ for 10 minutes followed by 20 minutes of stirring. Finally, Di-isopropylethylamine (15.1 mL, 86.7 mmol) and boron trifluoride diethyl etherate (15.3 mL, 124 mmol) were added after which the mixture was left stirring for 24 hours under N_2_. 2:3 hexane:ethyl acetate was added to the reaction medium after which dichloromethane was removed under educed pressure. After purification on silica (2:3 hexane:ethyl acetate) the product was isolated as a red, crystalline solid (1.07 g, 25% yield). 1H NMR (400 MHz, CDCl3) δ 7.52 (d, J = 8.1 Hz, 2H), 7.43 (d, J = 8.1 Hz, 2H), 6.69 (d, J = 4.1 Hz, 2H), 6.27 (d, J = 4.1 Hz, 2H), 4.46 (s, 2H), 2.65 (s, 6H). Step 3: Synthesis of 1,4,8,11-Tetra(prop-2-yn-1-yl)-1,4,8,11-tetraazacyclotetradecane based on (Counsell et al., 2016). To a solution of 1,4,8,11-tetraazacyclotetradecane (500 mg, 2.49 mmol) in 5 mL of acetonitrile 5 mL of 1M NaOH was added followed by 1.1 mL 80% propargyl bromide solution (in toluene, 10.2 mmol). The mixture was left overnight with very gentle stirring. The precipitate was collected, washed with hexane and dried under vacuum to yield the desired product (236 mg, 27% yield). 1H NMR (400 MHz, CDCl3) δ 3.44 (s, 8H), 2.62 (s, 16H), 2.17 (s, 4H), 1.77 (s, 4H), 1.61 (s, 4H). Step 4: Synthesis (2-azidoethyl)trimethylammonium bromide (based on (Francavilla et al., 2009)). To 30 mL of DMF (2-bromoethyl)trimethylammonium bromide (1 g, 4.05 mmol) and NaN_3_ (0.65 g, 10.1 mmol) were added. After stirring overnight the mixture is concentrated and diluted with THF after which the resulting precipitate was collected and dried to yield 2-azidoethyl)trimethylammonium bromide (480 mg, 57% yield). 1H NMR (400 MHz, D2O) δ 4.01 (s, 2H), 3.69 – 3.60 (m, 2H), 3.25 (s, 9H). Step 5: Synthesis of the cell wall mechano-probe (based on (Lipshutz and Taft, 2006)). To a 0.5 ml microwave vial N3-BODIPY (60 mg, 0.17 mmol), 1,4,8,11-Tetra(prop-2-yn-1-yl)-1,4,8,11-tetraazacyclotetradecane (60 mg, 0.17 mg) and Cu impregnated activated charcoal (17 mg) were combined. 0.4 mL 1,4-dioxane was added and the mixture was heated to 150 LC for 20 minutes under continuous stirring in a Biotage initiator+ system. After filtration (celite) the reaction mixture was dried. From the resulting solid 39 mg was added to a new 0.5 mL microwave vial after which (2-azidoethyl)trimethylammonium bromide (87 mg, 0.42 mmol) and Cu impregnated activated charcoal (17 mg) were added. 0.25 mL MilliQ water and 0.25 mL 1,4-dioxane were added and the mixture was heated to 150 LC for 20 minutes under continuous stirring in a Biotage initiator+ system. The resulting mixture was filtered over an 0.5 C18 SPE column. 6.6 mg of product was isolated as a red crystalline solid and was used without any additional purification.

### FLIM imaging and analysis

Five-day old seedlings of Col-0, *exad1-1* and *exad1-3* were treated with 100 mM NaCl for 48 h. Analysis were performed with a Leica TCS SP8 inverted scanning confocal microscope coupled with a Becker-Hickl SPC830 time-correlated single photon counting (TCSPC) module was used for FLIM image acquisition. A Leica TCS SP5 X pulsed white light laser with a repetition rate of 40 MHz and an excitation wavelength of 488 nm was used as a laser source. Imaging was performed with a 63× 1.2 NA water immersion objective with a 256×256 pixel resolution. A line scanning speed of 400 Hz was used, and the emission was collected between 500 nm and 550 nm onto a Leica HyD SMD hybrid photodetector.

### Statistical analysis

Statistical analysis in this study was performed in R. The normal distributions of phenotypic data were fitted in a linear model and two-way ANOVA was performed followed with contrasts post-hoc test. Treatments and genotypes were selected as the factors, and post-hoc contrasts comparison was performed with the ghlt (multcomp) package. For comparison between two groups, a T-test was performed.

## Acknowledgements

We thank Leónie Bentsink and Leo Willems of the Seed lab, Laboratory of Plant Physiology, Wageningen University & Research for providing HapMap seeds for the GWAS screen. We also thank David E. Salt and Paulina Flis from Future Food Beacon of Excellence and School of Biosciences, University of Nottingham, United Kingdom for ion measurements. Yutao Zou was sponsored by the China Scholarship Council (CSC), Christa Testerink acknowledges support from the European Research Council (ERC) through the European Union’s Horizon 2020 Research and Innovation program (ERC Consolidator Grant agreement 724321) and the Dutch Research Council (NWO) (Vici grant VI.C.192.033).

## Author Contributions

C.T. conceived the project, C.T., Yx.Z, Yut.Z and N.GB designed experiments and wrote the manuscript draft, which all other authors read and gave feedback on. C.T, Yx.Z and N.GB guided the research. Yut.Z performed the GWAS and the SITA assays with the help of Y.C., P.K. and J.A.D, and P.K., J.L. and J.A.D. were responsible for the halotropism experiments. P.K., N.GB and T.E performed the cell wall analysis and the dot-blot experiments. H.L. contributed to the gene expression analyses, T.P.N performed haplotype analysis and E.vZ. and. I.T.K provided scripts for root angle analysis. M.M.J performed and interpreted GWAS analysis. B.J and B.L.P performed CoMPP analysis. M.G, J.V, and T.K. guided and performed TEM imaging. M.B. and J.S performed and analyzed the cell wall mechano-probe experiments. J.L, T.dZ and P.K. contributed to the TEM and FLIM experiments.

## Competing Interests

The authors declare that there is no conflict of interest.

## Notes

### Competing Interest Statement

The authors have declared no competing interest.

### Summary of Updates

New data have been added, figures and text revised accordingly, and additional authors, who contributed to the revision, have been added.

## References

1. Alonso-Blanco, C. et al. (2016). 1,135 Genomes Reveal the Global Pattern of Polymorphism in Arabidopsis thaliana. Cell 166: 481–491.

2. Barberon, M., Vermeer, J.E.M., De Bellis, D., Wang, P., Naseer, S., Andersen, T.G., Humbel, B.M., Nawrath, C., Takano, J., Salt, D.E., and Geldner, N. (2016). Adaptation of Root Function by Nutrient-Induced Plasticity of Endodermal Differentiation. Cell 164: 447–459.

3. Baumberger, N., Ringli, C., and Keller, B. (2001). The chimeric leucine-rich repeat/extensin cell wall protein LRX1 is required for root hair morphogenesis in Arabidopsis thaliana. Genes Dev 15: 1128–1139.

4. Baumberger, N., Steiner, M., Ryser, U., Keller, B., and Ringli, C. (2003). Synergistic interaction of the two paralogous Arabidopsis genes LRX1 and LRX2 in cell wall formation during root hair development. Plant J 35: 71–81.

5. Butler, D.G., Cullis, B.R., Gilmour, A.R., and Gogel, B.J. ASReml-R reference manual.

6. Byrt, C.S., Munns, R., Burton, R.A., Gilliham, M., and Wege, S. (2018). Root cell wall solutions for crop plants in saline soils. Plant Science 269: 47–55.

7. Cannon, M.C., Terneus, K., Hall, Q., Tan, L., Wang, Y., Wegenhart, B.L., Chen, L., Lamport, D.T.A., Chen, Y., and Kieliszewski, M.J. (2008). Self-assembly of the plant cell wall requires an extensin scaffold. Proc Natl Acad Sci U S A 105: 2226–2231.

8. Castilleux, R., Plancot, B., Gügi, B., Attard, A., Loutelier-Bourhis, C., Lefranc, B., Nguema-Ona, E., Arkoun, M., Yvin, J.-C., Driouich, A., and Vicré, M. (2020). Extensin arabinosylation is involved in root response to elicitors and limits oomycete colonization. Ann Bot 125: 751– 763.

9. Chen, Y., Dong, W., Tan, L., Held, M.A., and Kieliszewski, M.J. (2015). Arabinosylation Plays a Crucial Role in Extensin Cross-linking In Vitro. Biochem Insights 8: 1–13.

10. Counsell, A.J., Jones, A.T., Todd, M.H., and Rutledge, P.J. (2016). A direct method for the N-tetraalkylation of azamacrocycles. Beilstein J. Org. Chem. 12: 2457–2461.

11. Deolu-Ajayi, A.O., Meyer, A.J., Haring, M.A., Julkowska, M.M., and Testerink, C. (2019). Genetic Loci Associated with Early Salt Stress Responses of Roots. iScience 21: 458–473.

12. Dinneny, J.R., Long, T.A., Wang, J.Y., Jung, J.W., Mace, D., Pointer, S., Barron, C., Brady, S.M., Schiefelbein, J., and Benfey, P.N. (2008). Cell identity mediates the response of Arabidopsis roots to abiotic stress. Science 320: 942–945.

13. Duan, A.-Q., Tao, J.-P., Jia, L.-L., Tan, G.-F., Liu, J.-X., Li, T., Chen, L.-Z., Su, X.-J., Feng, K., Xu, Z.-S., and Xiong, A.-S. (2020). AgNAC1, a celery transcription factor, related to regulation on lignin biosynthesis and salt tolerance. Genomics 112: 5254–5264.

14. Egelund, J., Obel, N., Ulvskov, P., Geshi, N., Pauly, M., Bacic, A., and Petersen, B.L. (2007). Molecular characterization of two Arabidopsis thaliana glycosyltransferase mutants, rra1 and rra2, which have a reduced residual arabinose content in a polymer tightly associated with the cellulosic wall residue. Plant Mol Biol 64: 439–451.

15. Endler, A., Kesten, C., Schneider, R., Zhang, Y., Ivakov, A., Froehlich, A., Funke, N., and Persson, S. (2015). A Mechanism for Sustained Cellulose Synthesis during Salt Stress. Cell 162: 1353–1364.

16. Fangel, J.U., Jones, C.Y., Ulvskov, P., Harholt, J., and Willats, W.G.T. (2021). Analytical implications of different methods for preparing plant cell wall material. Carbohydr Polym 261: 117866.

17. Feng, W. et al. (2018). The FERONIA Receptor Kinase Maintains Cell-Wall Integrity during Salt Stress through Ca ^2+^ Signaling. Current Biology 28: 666–675.e5.

18. Francavilla, C., Low, E., Nair, S., Kim, B., Shiau, T.P., Debabov, D., Celeri, C., Alvarez, N., Houchin, A., Xu, P., Najafi, R., and Jain, R. (2009). Quaternary ammonium N,N-dichloroamines as topical, antimicrobial agents. Bioorganic & Medicinal Chemistry Letters 19: 2731–2734.

19. Galvan-Ampudia, C.S., Julkowska, M.M., Darwish, E., Gandullo, J., Korver, R.A., Brunoud, G., Haring, M.A., Munnik, T., Vernoux, T., and Testerink, C. (2013). Halotropism is a response of plant roots to avoid a saline environment. Current Biology 23: 2044–2050.

20. Gigli-Bisceglia, N. and Testerink, C. (2021). Fighting salt or enemies: shared perception and signaling strategies. Curr Opin Plant Biol 64: 102120.

21. Gigli-Bisceglia, N., van Zelm, E., Huo, W., Lamers, J., and Testerink, C. (2022). Arabidopsis root responses to salinity depend on pectin modification and cell wall sensing. Development: dev.200363.

22. Gille, S., Hänsel, U., Ziemann, M., and Pauly, M. (2009). Identification of plant cell wall mutants by means of a forward chemical genetic approach using hydrolases. Proc Natl Acad Sci U S A 106: 14699–14704.

23. Hasegawa, P.M., Bressan, R.A., Zhu, J.-K., and Bohnert, H.J. (2000). PLANT CELLULAR AND MOLECULAR RESPONSES TO HIGH SALINITY. Annu Rev Plant Physiol Plant Mol Biol 51: 463–499.

24. Jacobowitz, J.R., Doyle, W.C., and Weng, J.-K. (2019). PRX9 and PRX40 Are Extensin Peroxidases Essential for Maintaining Tapetum and Microspore Cell Wall Integrity during Arabidopsis Anther Development. Plant Cell 31: 848–861.

25. Julkowska, M.M., Hoefsloot, H.C.J., Mol, S., Feron, R., de Boer, G.-J., Haring, M.A., and Testerink, C. (2014). Capturing Arabidopsis root architecture dynamics with ROOT-FIT reveals diversity in responses to salinity. Plant Physiol 166: 1387–1402.

26. Julkowska, M.M., Klei, K., Fokkens, L., Haring, M.A., Schranz, M.E., and Testerink, C. (2016). Natural variation in rosette size under salt stress conditions corresponds to developmental differences between Arabidopsis accessions and allelic variation in the LRR-KISS gene. J Exp Bot 67: 2127–2138.

27. Julkowska, M.M., Saade, S., Agarwal, G., Gao, G., Pailles, Y., Morton, M., Awlia, M., and Tester, M. (2019). MVApp—Multivariate Analysis Application for Streamlined Data Analysis and Curation. Plant Physiol. 180: 1261–1276.

28. Julkowska, M.M. and Testerink, C. (2015). Tuning plant signaling and growth to survive salt. Trends in Plant Science 20: 586–594.

29. Karlova, R., Boer, D., Hayes, S., and Testerink, C. (2021). Root plasticity under abiotic stress. Plant Physiol 187: 1057–1070.

30. Kassambara, A. and Mundt, F. (2020). factoextra: Extract and Visualize the Results of Multivariate Data Analyses.

31. Korte, A., Vilhjálmsson, B.J., Segura, V., Platt, A., Long, Q., and Nordborg, M. (2012). A mixed-model approach for genome-wide association studies of correlated traits in structured populations. Nat Genet 44: 1066–1071.

32. Korver, R.A., van den Berg, T., Meyer, A.J., Galvan-Ampudia, C.S., Ten Tusscher, K.H.W.J., and Testerink, C. (2020). Halotropism requires phospholipase Dζ1-mediated modulation of cellular polarity of auxin transport carriers. Plant Cell Environ 43: 143–158.

33. Lamport, D.T.A., Kieliszewski, M.J., Chen, Y., and Cannon, M.C. (2011). Role of the Extensin Superfamily in Primary Cell Wall Architecture1. Plant Physiol 156: 11–19.

34. Lamport, D.T.A. and Várnai, P. (2013). Periplasmic arabinogalactan glycoproteins act as a calcium capacitor that regulates plant growth and development. New Phytol 197: 58–64.

35. Leszczuk, A., Kalaitzis, P., Blazakis, K.N., and Zdunek, A. (2020). The role of arabinogalactan proteins (AGPs) in fruit ripening—a review. Hortic Res 7: 1–12.

36. Lipshutz, B.H. and Taft, B.R. (2006). Heterogeneous CopperLinLCharcoalLCatalyzed Click Chemistry. Angew Chem Int Ed 45: 8235–8238.

37. Liu, X., Wolfe, R., Welch, L.R., Domozych, D.S., Popper, Z.A., and Showalter, A.M. (2016). Bioinformatic Identification and Analysis of Extensins in the Plant Kingdom. PLoS One 11: e0150177.

38. Lobet, G., Pagès, L., and Draye, X. (2011). A novel image-analysis toolbox enabling quantitative analysis of root system architecture. Plant Physiol 157: 29–39.

39. Marzol, E., Borassi, C., Bringas, M., Sede, A., Rodríguez Garcia, D.R., Capece, L., and Estevez, J.M. (2018). Filling the Gaps to Solve the Extensin Puzzle. Mol Plant 11: 645–658.

40. Matsubayashi, Y. (2014). Posttranslationally modified small-peptide signals in plants. Annu Rev Plant Biol 65: 385–413.

41. Michels, L., Bronkhorst, J., Kasteel, M., de Jong, D., Albada, B., Ketelaar, T., Govers, F., and Sprakel, J. (2022). Molecular sensors reveal the mechano-chemical response of Phytophthora infestans walls and membranes to mechanical and chemical stress. Cell Surf 8: 100071.

42. Mishler-Elmore, J.W., Zhou, Y., Sukul, A., Oblak, M., Tan, L., Faik, A., and Held, M.A. (2021). Extensins: Self-Assembly, Crosslinking, and the Role of Peroxidases. Frontiers in Plant Science 12.

43. Moller, I., Sørensen, I., Bernal, A.J., Blaukopf, C., Lee, K., Øbro, J., Pettolino, F., Roberts, A., Mikkelsen, J.D., Knox, J.P., Bacic, A., and Willats, W.G.T. (2007). High-throughput mapping of cell-wall polymers within and between plants using novel microarrays. Plant J 50: 1118–1128.

44. Møller, S.R. et al. (2017). Identification and evolution of a plant cell wall specific glycoprotein glycosyl transferase, ExAD. Sci Rep 7: 45341.

45. Munns, R. and Tester, M. (2008). Mechanisms of salinity tolerance. Annu Rev Plant Biol 59: 651– 681.

46. Ogawa-Ohnishi, M., Matsushita, W., and Matsubayashi, Y. (2013). Identification of three hydroxyproline O-arabinosyltransferases in Arabidopsis thaliana. Nat Chem Biol 9: 726– 730.

47. Pattathil, S. et al. (2010). A comprehensive toolkit of plant cell wall glycan-directed monoclonal antibodies. Plant Physiol 153: 514–525.

48. Pérez García, M., Zhang, Y., Hayes, J., Salazar, A., Zabotina, O.A., and Hong, M. (2011). Structure and interactions of plant cell-wall polysaccharides by two– and three-dimensional magic-angle-spinning solid-state NMR. Biochemistry 50: 989–1000.

49. Petersen, B.L., MacAlister, C.A., and Ulvskov, P. (2021). Plant Protein O-Arabinosylation. Front Plant Sci 12: 645219.

50. Robin, A.H.K., Matthew, C., Uddin, M.J., and Bayazid, K.N. (2016). Salinity-induced reduction in root surface area and changes in major root and shoot traits at the phytomer level in wheat. J Exp Bot 67: 3719–3729.

51. Seren, Ü., Vilhjálmsson, B.J., Horton, M.W., Meng, D., Forai, P., Huang, Y.S., Long, Q., Segura, V., and Nordborg, M. (2012). GWAPP: a web application for genome-wide association mapping in Arabidopsis. Plant Cell 24: 4793–4805.

52. Smallwood, M., Beven, A., Donovan, N., Neill, S. j., Peart, J., Roberts, K., and Knox, J. p. (1994). Localization of cell wall proteins in relation to the developmental anatomy of the carrot root apex. The Plant Journal 5: 237–246.

53. Sun, F., Zhang, W., Hu, H., Li, B., Wang, Y., Zhao, Y., Li, K., Liu, M., and Li, X. (2008). Salt Modulates Gravity Signaling Pathway to Regulate Growth Direction of Primary Roots in Arabidopsis. Plant Physiol 146: 178–188.

54. Velasquez, S.M. et al. (2015). Low Sugar Is Not Always Good: Impact of Specific O-Glycan Defects on Tip Growth in Arabidopsis1. Plant Physiol 168: 808–813.

55. Velasquez, S.M. et al. (2011). O-glycosylated cell wall proteins are essential in root hair growth. Science 332: 1401–1403.

56. Voxeur, A., Wang, Y., and Sibout, R. (2015). Lignification: different mechanisms for a versatile polymer. Curr Opin Plant Biol 23: 83–90.

57. Weigel, D. and Mott, R. (2009). The 1001 genomes project for Arabidopsis thaliana. Genome Biol 10: 107.

58. Xie, D., Ma, L., Samaj, J., and Xu, C. (2011). Immunohistochemical analysis of cell wall hydroxyproline-rich glycoproteins in the roots of resistant and susceptible wax gourd cultivars in response to Fusarium oxysporum f. sp. Benincasae infection and fusaric acid treatment. Plant Cell Rep 30: 1555–1569.

59. Xu, C., Zhao, L., Pan, X., and Samaj, J. (2011). Developmental localization and methylesterification of pectin epitopes during somatic embryogenesis of banana (Musa spp. AAA). PLoS One 6: e22992.

60. Yan, J., Liu, Y., Yang, L., He, H., Huang, Y., Fang, L., Scheller, H.V., Jiang, M., and Zhang, A. (2021). Cell wall β-1,4-galactan regulated by the BPC1/BPC2-GALS1 module aggravates salt sensitivity in Arabidopsis thaliana. Mol Plant 14: 411–425.

61. Yeats, T., Vellosillo, T., Sorek, N., IbLLez, A., and Bauer, S. (2016). Rapid Determination of Cellulose, Neutral Sugars, and Uronic Acids from Plant Cell Walls by One-step Two-step Hydrolysis and HPAEC-PAD. BIO-PROTOCOL 6.

62. van Zelm, E., Zhang, Y., and Testerink, C. (2020). Salt Tolerance Mechanisms of Plants. Annual Review of Plant Biology 71: 403–433.

63. Zhao, C., Zayed, O., Yu, Z., Jiang, W., Zhu, P., Hsu, C.C., Zhang, L., Andy Tao, W., Lozano-Durán, R., and Zhu, J.K. (2018). Leucine-rich repeat extensin proteins regulate plant salt tolerance in Arabidopsis. Proceedings of the National Academy of Sciences of the United States of America 115: 13123–13128.

64. Zhao, C., Zayed, O., Zeng, F., Liu, C., Zhang, L., Zhu, P., Hsu, C.-C., Tuncil, Y.E., Tao, W.A., Carpita, N.C., and Zhu, J.-K. (2019). Arabinose biosynthesis is critical for salt stress tolerance in Arabidopsis. New Phytol 224: 274–290.

65. Zou, Y., Zhang, Y., and Testerink, C. (2022). Root dynamic growth strategies in response to salinity. Plant Cell Environ 45: 695–704.

