## Supplementary material for "Cell wall extensin arabinosylation is required for root directional response to salinity": Suppl Figures

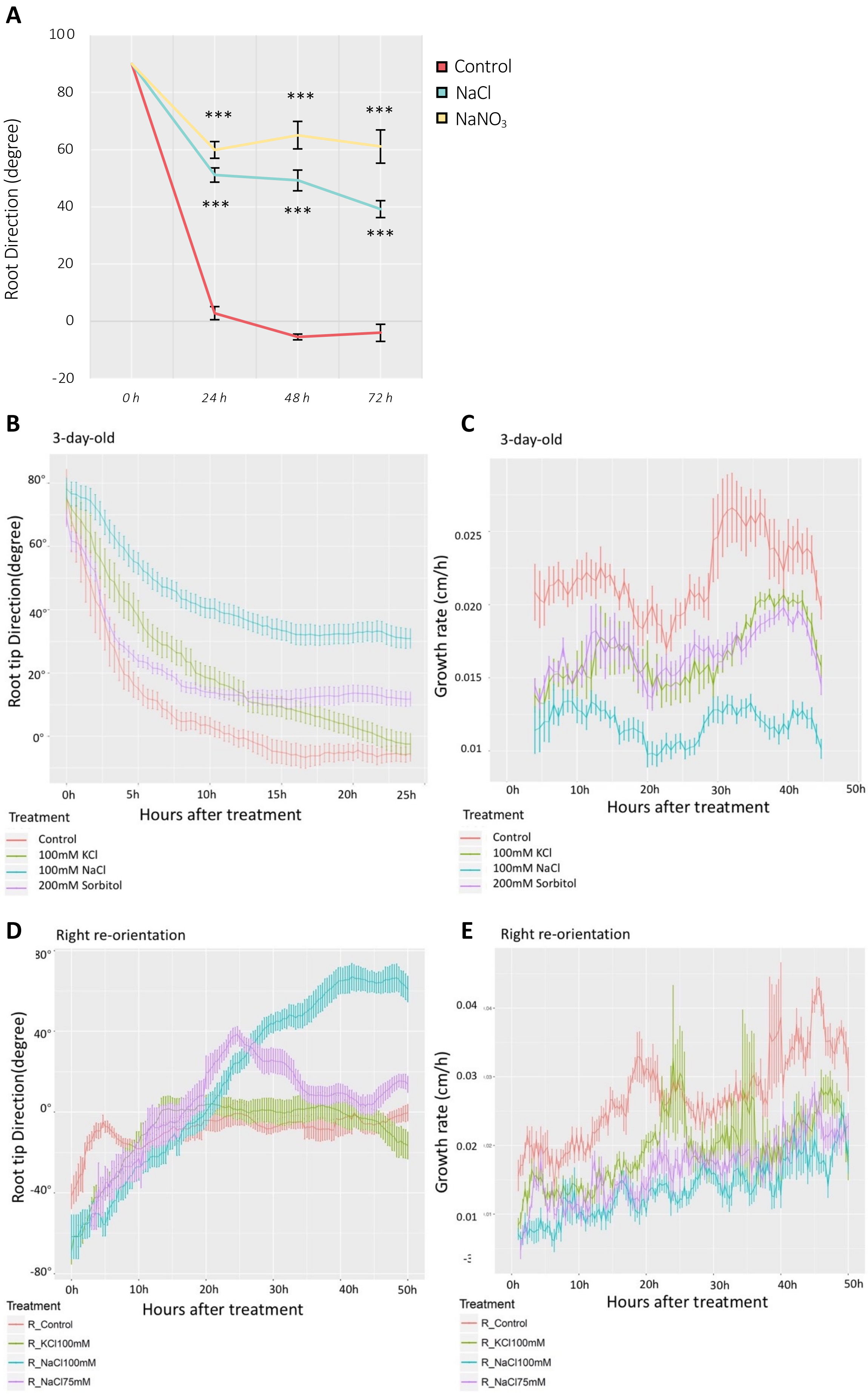


**Figure S1**. Root Tip direction (RTD) in time-lapse SITA is modulated by NaCl treatment. **A** Root angle was analyzed in SITA in Col-0 seedlings treated with (control, 100 mM NaCl, 100 mM $NaNO_3$) at 24, 48 and 72 h. Comparisons between control and NaCl or control and $NaNO_3$ have been performed by using multiple paired T-tests coupled with Benjamini and Hochberg False Discovery Rate correction. ***, P < 0.001. **B** and **C**, quantification of RTD **(B)** and growth rate **(C)** of 3-day-old Col-0 seedlings under different treatments (control, 100 mM NaCl, 100 mM KCl and 200 mM sorbitol) over 24 h. Values represent means ± SE of 40 seedlings from 2 plates. **D** and **E**, quantification of RTD **(E)** and growth rate **(E)** of 3-day-old Col-0 seedlings under different treatments (control, 100 mM NaCl, 100 mM KCl and 200 mM Sorbitol) over 24 h. Values represent means ± SE of 35 seedlings from 1 plate. C and D are from 2 independent experiments.

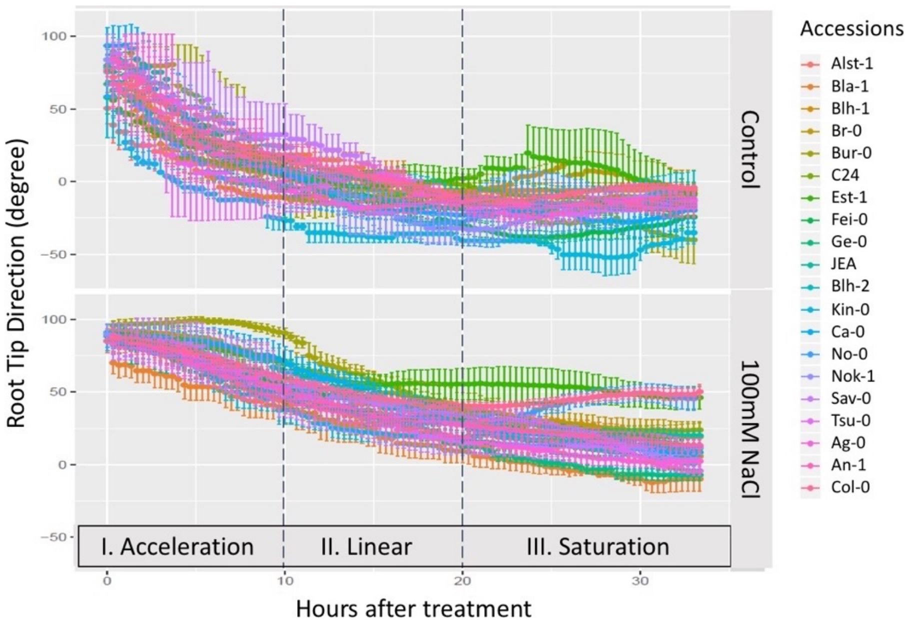


**Figure S2.** Natural variation observed in root tip direction (RTD) of 20 Arabidopsis accessions analyzed with the SITA time-lapse system. Four-day-old seedlings of 20 Arabidopsis accessions were transferred to agar plates with or without 100 mM NaCl. The different colors represent the analyzed accessions. Dynamic RTD responses were followed in three phases that are indicated with dash lines: I) Acceleration, II) Linear and III) Saturation. Values represent means ± SE of 5 seedlings. Data are representative of two independent experiments.

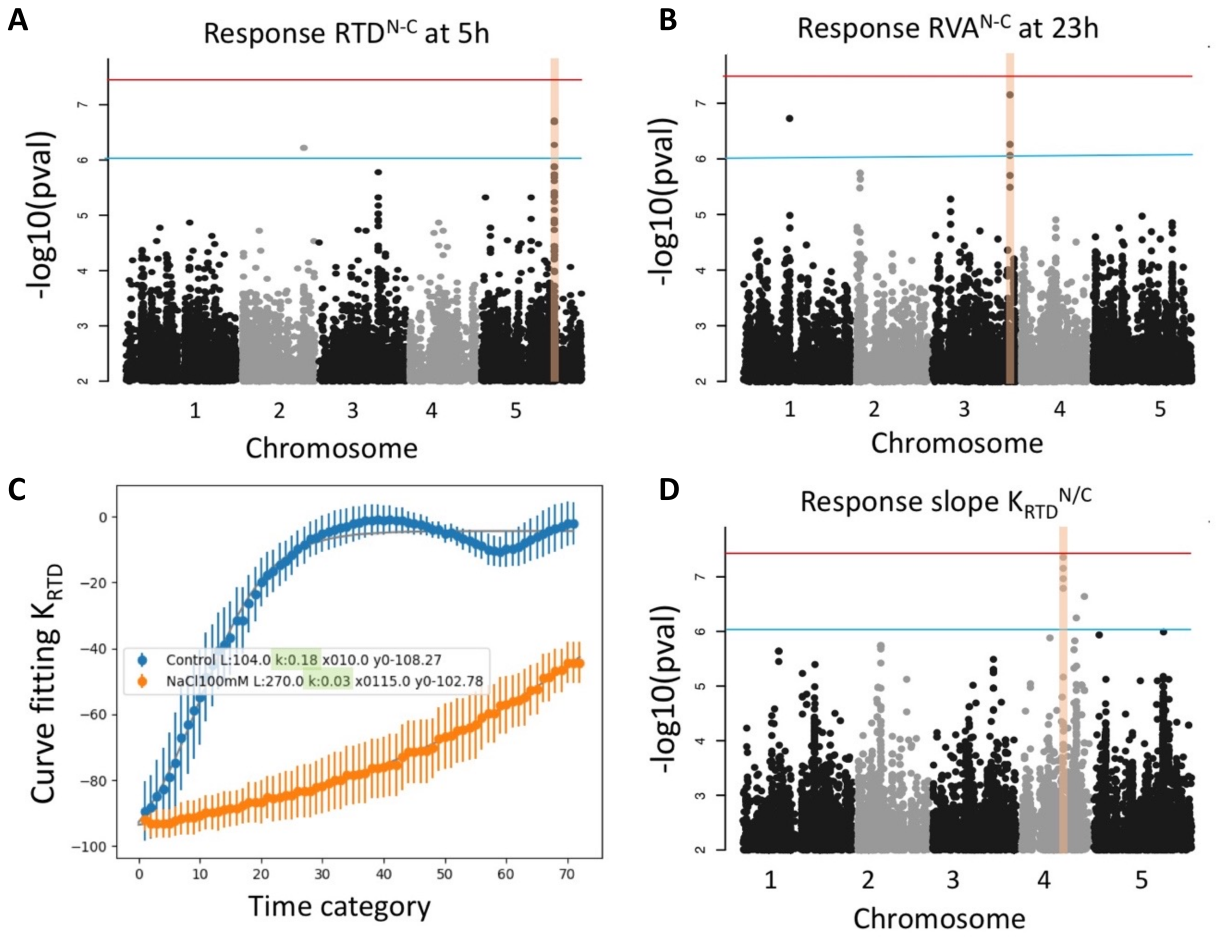


**Figure S3. Candidate loci for different traits in SITA mapped by GWAS using 345 Arabidopsis accessions.** Manhattan plots illustrating the association of single nucleotide polymorphisms (SNPs) with two root response traits, namely, Response Root Tip Direction at 5 hours ($RTD^{N\text{-}C}$) in panel **A** and Response Root Vector Angle at 23 hours ($RAV^{N\text{-}C}$) in panel **B,** $RTD^{N\text{-}C}$ and $RAV^{N\text{-}C}$ denote the differences in Root Tip Directions (RTDs) and Root Vector Angles (RAVs), respectively, between salt-stressed and control conditions. **C,** depicts the calculated $K_{RTD}$ (Kinetic Relative Root Tip Direction) for the Col-0 accession, derived from a 4-parameter logistic regression (4PL) model fit to relative RTD data across various accessions, describing the rate of relative root response under salt stress. Python was employed for parameter calculation through curve fitting (script available upon request). The black boxes highlight the 'k' values, representing the KRTD parameter, for both control and NaCl treatments as determined by the model. **D,** Manhattan plot for the SNPs associated with $K_{RTD}{}^{N/C}$. $K^{N/C}$ is the response in fitted rate of exponential decay of RTDs, $K_{RTD}{}^{N/C}$ represents $K_{RTD}$ under salt condition divided by that under control condition. In each Manhattan plot, different chromosomes are indicated by gray or black color as X-axis; the Y-axis shows LOD score. All *loci* above the arbitrary threshold of LOD=6.0 (bule line) were highlighted in orange and listed in Table S2. Bonferroni threshold = 7.55 (red line) was determined by the $-\log_{10}$(p-value / # SNPs) for SNPs with minor allele frequency > 0.05.

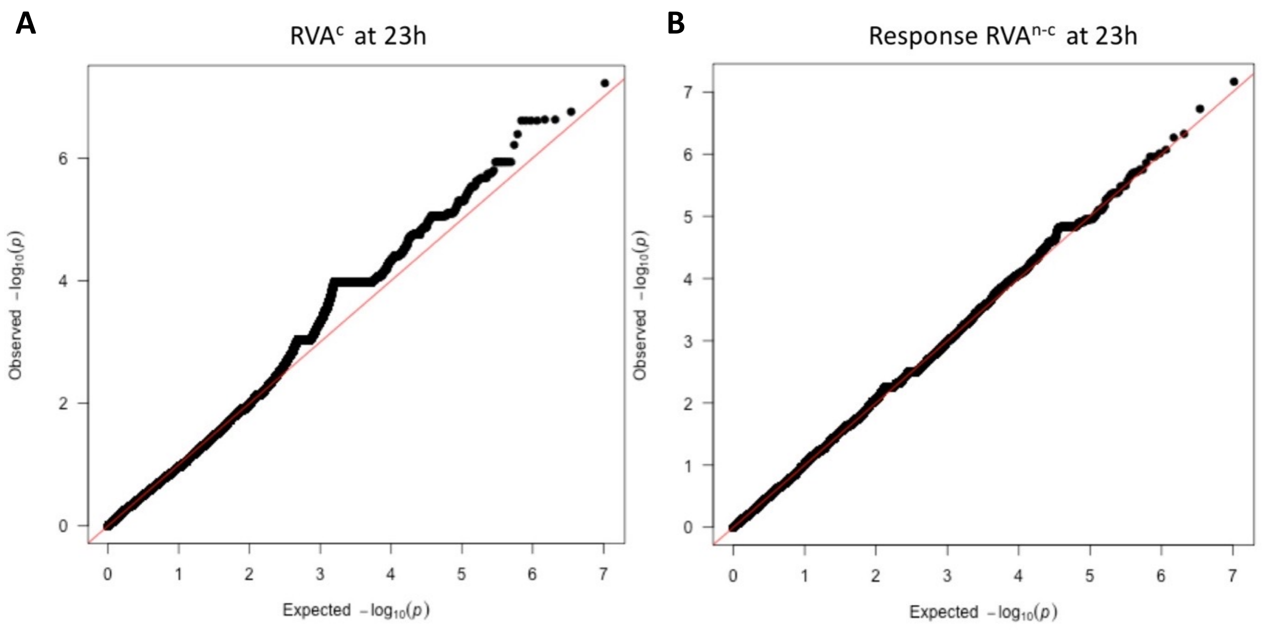


**Figure S4. QQ-plots for traits in SITA mapped by GWAS using 345 Arabidopsis accessions.** QQ-plots for root vector angle under control condition ($RAV^{C}$) **(A)** and root response root vector angle ($RAV^{N\text{-}C}$) at 23 h **(B)**.

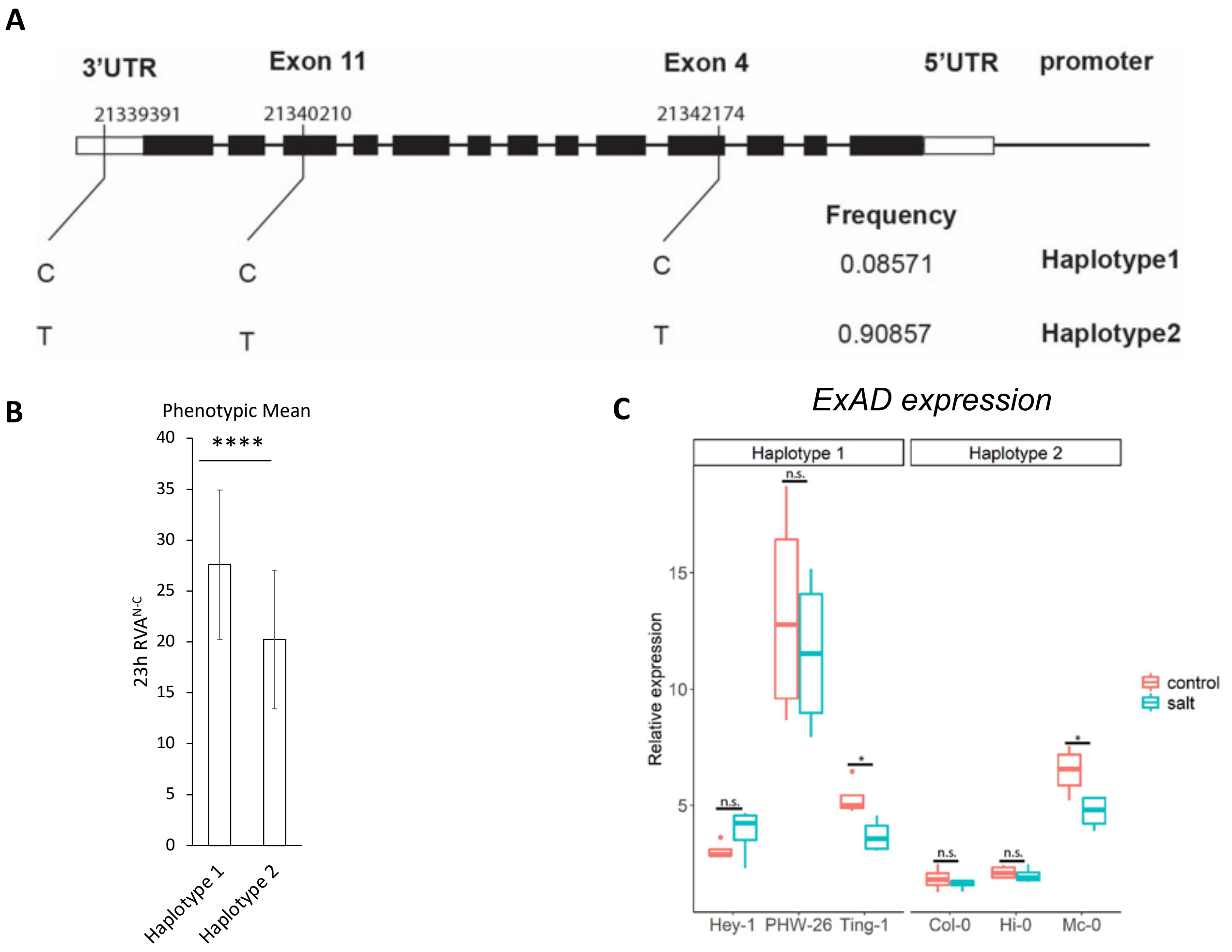


**Figure S5. Natural variation analysis of ExAD gene. A**, Two haplotypes were identified for ExAD (*AT3G57630*) using 3 significant SNPs found in GWAS (SNP at position 21339391, 21340210 and 21342174) based on genomic variations and haplotype frequency in Arabidopsis Hapmap accessions. **B**, Average of root vector angle values ($RVA^{N-C}$) at 23 h under salt condition subtracted by RVA under control condition in two haplotypes of ExAD. Data represent means ± SD. **C**, Expression of ExAD under salt (100 mM NaCl for 48 h) and control conditions in tested accessions from two haplotypes relative to housekeeping gene (*AT2G43770*). Data represent means ± SE. Statistical analyses in (**B)** and (**C)** were determined using Student's T-Test. F-test was performed to check equal variances of samples. Significant differences were determined by Two-sample T-Test with equal variance otherwise Welch's t-test (*, $p<0.05$; ****,$p<0.001$; $p>0.05$, n.s.).

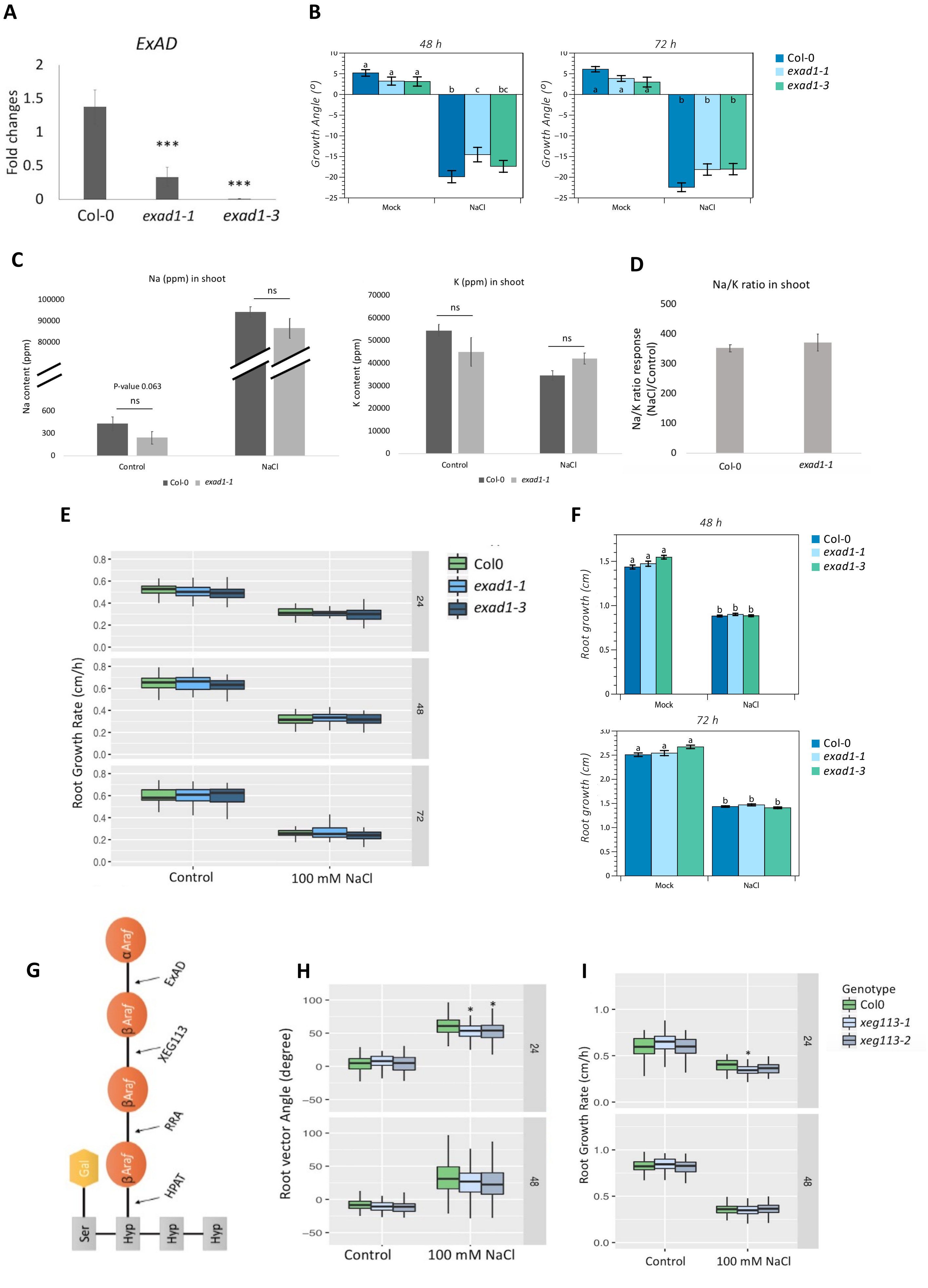
A
ExAD
Fold changes
2
1.5
1
0.5
0
***
***
Col-0
exad1-1
exad1-3
B
48 h
72 h
Growth Angle (º)
a
b
c
bc
Mock
NaCl
Col-0
exad1-1
exad1-3
C
Na (ppm) in shoot
Na content (ppm)
100000
90000
80000
600
300
0
P-value 0.063
ns
Control
NaCl
Col-0
exad1-1
K (ppm) in shoot
K content (ppm)
70000
60000
50000
40000
30000
20000
10000
0
D
Na/K ratio in shoot
Na/K ratio response (NaCl/Control)
500
400
300
200
100
0
Col-0
exad1-1
E
Root Growth Rate (cm/h)
Col0
exad1-1
exad1-3
24
48
72
Control
100 mM NaCl
F
48 h
72 h
Root growth (cm)
Mock
NaCl
Col-0
exad1-1
exad1-3
G
αAraf
βAraf
βAraf
βAraf
Gal
ExAD
XEG113
RRA
HPAT
Ser
Hyp
Hyp
Hyp
H
Root vector Angle (degree)
24
48
Control
100 mM NaCl
I
Root Growth Rate (cm/h)
Genotype
Col0
xeg113-1
xeg113-2
24
48
Control
100 mM NaCl

**Figure S6. A.** Expression of *ExAD* in 8-day-old *Arabidopsis* seedlings of Col-0, *exad1-1* and *exad1-3* mutants measured through qRT-PCR. Primers used for the analysis are reported in Table S4. Values represent the average ± SE of nine biological replicates, each containing at least 5 seedlings. Statistical analysis was done with Student's T-test. **B**, Absolute root angle values of experiment in Figure 3C was performed as in (Galvan-Ampudia et al., 2013) in 5-day old Col-0, *exad1-1* and *exad1-3* seedlings. Seedlings were transferred to agar plates with/without NaCl gradient and root growth in cm was measured after 48 and 72 h from the transferring time point. Values represent means ± SE from 48 seedlings Seedlings were treated for 48 and 72 h and root bending analysed after plate scanning with Smartroot (Fiji Plugin). Non-parametric analysis (Shapiro-Wilk test, $p < 0.05$) was performed. Kruskal-Wallis test was used to detect group differences, and Dunn's test determined specific significant differences ($p < 0.05$). **C,** Quantification of $Na^+$ content (ppm) and $K^+$ content (ppm), and Na+/K+ ratio (NaCl treatment/Control treatment) (**D)** was performed in shoots of Col-0 and the *exad1-1* mutant. Three-week-old plants were grown hydroponically and transferred to liquid media containing either NaCl (150 mM) or control (0 mM) for 4 days. The shoots were harvested for ion measurement, and data were normalized by fresh weight. Values represent means ± SE from 4 biological replicates, each containing 2 shoots from independent plants. Statistical analysis was performed using Student's T-test. No significant differences (ns) were detected. **E,** Quantification of root growth rate of Col-0, *exad1-1* and *exad1-3* mutants in SITA. Four-day-old seedlings were transferred to agar plates with or without 100 mM NaCl for 24, 48 and 72 h. Values represent means ± SE from 50 seedlings from 10 plates, each plate containing 5 seedlings. Data are representative of two independent experiments. Statistical analysis was performed by using two-way ANOVA with contrasts post-hoc. **F,** Quantification of root growth rate of Col-0, *exad1-1* and *exad1-3* mutants, in halotropism (Figure 3C) is expressed in cm. Treatment and statistical analysis were performed as in (B). **G,** Schematic diagram of extensin repeat side-chain Hyp-$Araf_{1\text{-}4}$ and extensin arabinosylation enzymes. Serine is substituted with galactose and the hydroxyprolines (Hyps) are substituted with β- arabinofuranoses (β-Araf) and α-arabinofuranoses (α-Araf). HYDROXYPROLINE ARABINOSYLTRANSFERASES 1-3 (HPAT1-3) adds the first arabinose (Hyp-$Araf_1$) to Hydroxyprolines (Hyps) (Ogawa-Ohnishi et al., 2013). REDUCED RESIDUAL ARABINOSE 1-3 (RRA1-3) adds the second arabinose residue (Hyp-$Araf_2$) (Egelund et al., 2007; Velasquez et al., 2011). XYLOGLUCAN ENDOGLUCANASE 113 (XEG113) adds the third arabinose residue (Hyp-$Araf_3$) (Gille et al., 2009). EXTENSIN DEFICIENT ARABINOSE (ExAD), ExAD adds the fourth arabinose residue (Hyp-$Araf_4$) on the extensin repeat **H,** Quantification of root vector angle of Col0, *xeg113-1* and *xeg113-2* mutants in SITA. Four-day-old seedlings transferred to agar plate with or without 100 mM NaCl for 24 and 48 h. Values represent means ± SE from 7 biological replicates (plates) and 10 technical replicates (seedlings). Error bars represent SE of the mean. Statistical analysis was done using two-way ANOVA with contrasts post-hoc. Asterisks indicate statistically significant differences compared to Col-0 (p-values ***P< 0.001; **P < 0.01; *P < 0.05). **I,** Quantification of root growth rate of Col-0, *xeg113-1* and *xeg113-2* mutants in SITA. Four-day-old seedlings transferred to agar plate with or without 100 mM NaCl for 24 and 48 h. Values represent the means ± SE of 7 biological replicates (plates) and 10 technical replicates (seedlings). Statistical analysis was done using two-way ANOVA with contrasts post-hoc. Asterisks indicate statistically significant differences compared to Col-0 (***P < 0.001; **P < 0.01; *P < 0.05).

| Extraction | Geno types | Treatment | JIM5 | JIM7 | LM5 | LM6 | LM7 | LM8 | LM13 | LM16 | LM18 | LM19 | LM20 | Inra RU1 | Inra RU2 | LM15 | LM25 | LM10 | LM11 | LM28 | LM23 | BS400-4 | LM21 | LM22 | CBM3a | LM1 | LM3 | JIM11 | JIM12 | JIM19 | JIM20 | LM2 | LM14 | JIM4 | JIM8 | JIM13 | JIM14 | JIM15 | JIM16 | JIM17 | MAC207 | anti-rat | anti-mouse | anti-his |
|---|---|---|---|---|---|---|---|---|---|---|---|---|---|---|---|---|---|---|---|---|---|---|---|---|---|---|---|---|---|---|---|---|---|---|---|---|---|---|---|---|---|---|---|---|
| CDTA Fraction | Col-0 | Control (0mM) | 68 | 100 | 0 | 24 | 0 | 0 | 0 | 0 | 57 | 76 | 86 | 45 | 59 | 8 | 12 | 0 | 0 | 0 | 0 | 0 | 0 | 0 | 0 | 0 | 69 | 39 | 37 | 0 | 32 | 31 | 43 | 0 | 0 | 16 | 0 | 0 | 21 | 90 | 20 | 0 | 0 | 0 |
| | Col-0 | Control (0mM) | 46 | 75 | 0 | 25 | 0 | 0 | 0 | 0 | 40 | 55 | 71 | 37 | 48 | 9 | 13 | 0 | 0 | 0 | 0 | 0 | 0 | 0 | 0 | 0 | 64 | 35 | 44 | 0 | 35 | 29 | 40 | 0 | 0 | 17 | 0 | 0 | 19 | 65 | 23 | 0 | 0 | 0 |
| | Col-0 | Control (0mM) | 55 | 98 | 0 | 20 | 0 | 0 | 0 | 0 | 48 | 68 | 95 | 41 | 54 | 6 | 10 | 0 | 0 | 0 | 0 | 0 | 0 | 0 | 0 | 0 | 48 | 32 | 46 | 0 | 33 | 30 | 39 | 0 | 0 | 13 | 0 | 0 | 13 | 89 | 20 | 0 | 0 | 0 |
| | *exad1-1* | Control (0mM) | 29 | 25 | 0 | 14 | 0 | 0 | 0 | 0 | 49 | 67 | 0 | 32 | 37 | 11 | 9 | 0 | 0 | 0 | 0 | 0 | 0 | 0 | 0 | 0 | 37 | 0 | 34 | 0 | 27 | 20 | 26 | 0 | 0 | 9 | 0 | 0 | 9 | 29 | 13 | 0 | 0 | 0 |
| | *exad1-1* | Control (0mM) | 30 | 26 | 0 | 9 | 0 | 0 | 0 | 0 | 50 | 71 | 0 | 25 | 26 | 8 | 7 | 0 | 0 | 0 | 0 | 0 | 0 | 0 | 0 | 0 | 30 | 0 | 33 | 0 | 30 | 17 | 20 | 0 | 0 | 6 | 0 | 0 | 6 | 27 | 8 | 0 | 0 | 0 |
| | *exad1-1* | Control (0mM) | 30 | 30 | 0 | 13 | 0 | 0 | 0 | 0 | 52 | 71 | 0 | 34 | 36 | 9 | 10 | 0 | 0 | 0 | 0 | 0 | 0 | 0 | 0 | 0 | 34 | 0 | 36 | 0 | 28 | 18 | 19 | 0 | 0 | 7 | 0 | 0 | 10 | 30 | 9 | 0 | 0 | 0 |
| | *exad1-3* | Control (0mM) | 58 | 72 | 0 | 17 | 0 | 0 | 0 | 0 | 63 | 83 | 45 | 34 | 39 | 7 | 9 | 0 | 0 | 0 | 0 | 0 | 0 | 0 | 0 | 0 | 41 | 0 | 41 | 0 | 33 | 20 | 20 | 0 | 0 | 5 | 0 | 0 | 9 | 62 | 8 | 0 | 0 | 0 |
| | *exad1-3* | Control (0mM) | 56 | 73 | 0 | 28 | 0 | 0 | 0 | 0 | 53 | 72 | 52 | 38 | 48 | 9 | 14 | 0 | 0 | 0 | 0 | 0 | 0 | 0 | 0 | 0 | 83 | 0 | 52 | 0 | 42 | 32 | 36 | 0 | 0 | 13 | 0 | 0 | 22 | 66 | 20 | 0 | 0 | 0 |
| | *exad1-3* | Control (0mM) | 71 | 98 | 0 | 18 | 0 | 0 | 0 | 0 | 56 | 74 | 86 | 36 | 42 | 10 | 12 | 0 | 0 | 0 | 0 | 0 | 0 | 0 | 0 | 0 | 57 | 0 | 50 | 0 | 39 | 28 | 34 | 0 | 0 | 11 | 0 | 0 | 12 | 86 | 16 | 0 | 0 | 0 |
| | Col-0 | NaCl (100mM) | 41 | 40 | 0 | 29 | 0 | 0 | 0 | 0 | 47 | 72 | 0 | 27 | 34 | 0 | 8 | 0 | 0 | 7 | 0 | 0 | 0 | 0 | 0 | 0 | 48 | 29 | 30 | 0 | 28 | 25 | 31 | 0 | 0 | 8 | 0 | 0 | 19 | 34 | 23 | 0 | 0 | 0 |
| | Col-0 | NaCl (100mM) | 35 | 39 | 0 | 25 | 0 | 0 | 0 | 0 | 42 | 62 | 0 | 23 | 25 | 6 | 7 | 0 | 0 | 5 | 0 | 0 | 0 | 0 | 0 | 0 | 43 | 35 | 29 | 0 | 33 | 24 | 30 | 0 | 0 | 9 | 0 | 0 | 21 | 35 | 23 | 0 | 0 | 0 |
| | Col-0 | NaCl (100mM) | 32 | 40 | 0 | 29 | 0 | 0 | 0 | 0 | 43 | 58 | 0 | 21 | 24 | 5 | 7 | 0 | 0 | 6 | 0 | 0 | 0 | 0 | 0 | 0 | 59 | 45 | 36 | 0 | 43 | 29 | 36 | 0 | 0 | 12 | 0 | 0 | 25 | 35 | 29 | 0 | 0 | 0 |
| | *exad1-1* | NaCl (100mM) | 41 | 57 | 0 | 25 | 0 | 0 | 0 | 0 | 43 | 64 | 29 | 18 | 15 | 6 | 6 | 0 | 0 | 0 | 0 | 0 | 0 | 0 | 0 | 0 | 58 | 0 | 37 | 0 | 36 | 24 | 25 | 0 | 0 | 0 | 0 | 0 | 13 | 53 | 19 | 0 | 0 | 0 |
| | *exad1-1* | NaCl (100mM) | 36 | 46 | 0 | 27 | 0 | 0 | 0 | 0 | 45 | 61 | 17 | 17 | 15 | 6 | 6 | 0 | 0 | 0 | 0 | 0 | 0 | 0 | 0 | 0 | 57 | 0 | 30 | 0 | 31 | 22 | 25 | 0 | 0 | 0 | 0 | 0 | 11 | 40 | 19 | 0 | 0 | 0 |
| | *exad1-1* | NaCl (100mM) | 49 | 65 | 0 | 29 | 0 | 0 | 0 | 0 | 56 | 76 | 40 | 24 | 23 | 5 | 8 | 0 | 0 | 0 | 0 | 0 | 0 | 0 | 0 | 0 | 61 | 0 | 33 | 0 | 32 | 23 | 28 | 0 | 0 | 0 | 0 | 0 | 12 | 50 | 22 | 0 | 0 | 0 |
| | *exad1-3* | NaCl (100mM) | 34 | 27 | 0 | 29 | 0 | 0 | 0 | 0 | 56 | 69 | 0 | 18 | 13 | 7 | 7 | 0 | 0 | 0 | 0 | 0 | 0 | 0 | 0 | 0 | 57 | 0 | 33 | 0 | 32 | 25 | 27 | 0 | 0 | 6 | 0 | 0 | 15 | 26 | 21 | 0 | 0 | 0 |
| | *exad1-3* | NaCl (100mM) | 34 | 28 | 0 | 31 | 0 | 0 | 0 | 0 | 55 | 69 | 0 | 12 | 9 | 6 | 6 | 0 | 0 | 0 | 0 | 0 | 0 | 0 | 0 | 0 | 62 | 0 | 35 | 0 | 37 | 28 | 28 | 0 | 0 | 6 | 0 | 0 | 15 | 24 | 22 | 0 | 0 | 0 |
| | *exad1-3* | NaCl (100mM) | 39 | 36 | 0 | 38 | 0 | 0 | 0 | 0 | 62 | 76 | 0 | 24 | 27 | 8 | 10 | 0 | 0 | 8 | 0 | 0 | 0 | 0 | 0 | 0 | 91 | 0 | 50 | 0 | 46 | 36 | 36 | 0 | 0 | 11 | 0 | 0 | 29 | 41 | 33 | 0 | 0 | 0 |
| NaOH-Fraction | Col-0 | Control (0mM) | 0 | 0 | 0 | 0 | 0 | 0 | 0 | 0 | 0 | 16 | 0 | 8 | 10 | 44 | 36 | 0 | 0 | 19 | 0 | 0 | 9 | 0 | 0 | 0 | 60 | 35 | 35 | 0 | 28 | 12 | 21 | 0 | 0 | 0 | 0 | 0 | 5 | 0 | 6 | 0 | 0 | 0 |
| | Col-0 | Control (0mM) | 0 | 0 | 0 | 0 | 0 | 0 | 0 | 0 | 0 | 15 | 0 | 8 | 9 | 28 | 30 | 0 | 0 | 16 | 0 | 0 | 0 | 0 | 0 | 0 | 67 | 33 | 37 | 0 | 34 | 15 | 19 | 0 | 0 | 0 | 0 | 0 | 9 | 0 | 5 | 0 | 0 | 0 |
| | Col-0 | Control (0mM) | 0 | 0 | 0 | 0 | 0 | 0 | 0 | 0 | 0 | 16 | 0 | 8 | 8 | 31 | 37 | 0 | 0 | 16 | 0 | 0 | 0 | 0 | 0 | 0 | 66 | 30 | 36 | 0 | 29 | 14 | 11 | 0 | 0 | 0 | 0 | 0 | 8 | 0 | 0 | 0 | 0 | 0 |
| | *exad1-1* | Control (0mM) | 0 | 0 | 0 | 0 | 0 | 0 | 0 | 0 | 0 | 12 | 0 | 0 | 0 | 39 | 32 | 0 | 0 | 16 | 0 | 0 | 0 | 0 | 0 | 0 | 36 | 26 | 27 | 0 | 31 | 14 | 11 | 0 | 0 | 0 | 0 | 0 | 13 | 0 | 0 | 0 | 0 | 0 |
| | *exad1-1* | Control (0mM) | 0 | 0 | 0 | 0 | 0 | 0 | 0 | 0 | 0 | 11 | 0 | 0 | 0 | 45 | 33 | 0 | 0 | 14 | 0 | 0 | 0 | 0 | 0 | 0 | 21 | 0 | 19 | 0 | 17 | 6 | 7 | 0 | 0 | 0 | 0 | 0 | 0 | 0 | 0 | 0 | 0 | 0 |
| | *exad1-1* | Control (0mM) | 0 | 0 | 0 | 0 | 0 | 0 | 0 | 0 | 0 | 12 | 0 | 0 | 0 | 32 | 26 | 0 | 0 | 16 | 0 | 0 | 0 | 0 | 0 | 0 | 37 | 0 | 29 | 0 | 26 | 10 | 12 | 0 | 0 | 0 | 0 | 0 | 0 | 0 | 0 | 0 | 0 | 0 |
| | *exad1-3* | Control (0mM) | 0 | 0 | 0 | 0 | 0 | 0 | 0 | 0 | 0 | 10 | 0 | 0 | 0 | 34 | 29 | 0 | 0 | 12 | 0 | 0 | 0 | 0 | 0 | 0 | 25 | 0 | 24 | 0 | 22 | 9 | 10 | 0 | 0 | 0 | 0 | 0 | 0 | 0 | 0 | 0 | 0 | 0 |
| | *exad1-3* | Control (0mM) | 0 | 0 | 0 | 0 | 0 | 0 | 0 | 0 | 0 | 17 | 0 | 7 | 0 | 47 | 38 | 0 | 28 | 28 | 0 | 0 | 7 | 0 | 0 | 0 | 32 | 0 | 23 | 0 | 17 | 6 | 6 | 0 | 0 | 0 | 0 | 0 | 0 | 0 | 0 | 0 | 0 | 0 |
| | *exad1-3* | Control (0mM) | 0 | 0 | 0 | 0 | 0 | 0 | 0 | 0 | 0 | 14 | 0 | 0 | 0 | 52 | 45 | 0 | 0 | 14 | 0 | 0 | 0 | 0 | 0 | 0 | 38 | 0 | 29 | 0 | 26 | 10 | 8 | 0 | 0 | 0 | 0 | 0 | 0 | 0 | 0 | 0 | 0 | 0 |
| | Col-0 | NaCl (100mM) | 0 | 0 | 0 | 6 | 0 | 0 | 0 | 0 | 0 | 20 | 0 | 6 | 5 | 30 | 37 | 0 | 6 | 26 | 0 | 0 | 0 | 0 | 0 | 0 | 51 | 11 | 30 | 0 | 0 | 0 | 0 | 0 | 0 | 0 | 0 | 0 | 0 | 0 | 0 | 0 | 0 | 0 |
| | Col-0 | NaCl (100mM) | 0 | 0 | 0 | 5 | 0 | 0 | 0 | 0 | 0 | 19 | 0 | 6 | 0 | 33 | 36 | 0 | 7 | 26 | 0 | 0 | 0 | 0 | 0 | 0 | 55 | 14 | 31 | 0 | 0 | 0 | 0 | 0 | 0 | 0 | 0 | 0 | 0 | 0 | 0 | 0 | 0 | 0 |
| | Col-0 | NaCl (100mM) | 0 | 0 | 0 | 5 | 0 | 0 | 0 | 0 | 0 | 18 | 0 | 6 | 6 | 35 | 36 | 0 | 0 | 26 | 0 | 0 | 0 | 0 | 0 | 0 | 54 | 11 | 32 | 0 | 0 | 0 | 0 | 0 | 0 | 0 | 0 | 0 | 0 | 0 | 0 | 0 | 0 | 0 |
| | *exad1-1* | NaCl (100mM) | 0 | 0 | 0 | 9 | 0 | 0 | 0 | 0 | 0 | 15 | 0 | 0 | 0 | 51 | 40 | 0 | 12 | 28 | 0 | 0 | 0 | 0 | 0 | 0 | 21 | 0 | 17 | 0 | 9 | 0 | 0 | 0 | 0 | 0 | 0 | 0 | 0 | 0 | 0 | 0 | 0 | 0 |
| | *exad1-1* | NaCl (100mM) | 0 | 0 | 0 | 9 | 0 | 0 | 0 | 0 | 0 | 15 | 0 | 0 | 0 | 51 | 42 | 0 | 6 | 26 | 0 | 0 | 0 | 0 | 0 | 0 | 21 | 0 | 18 | 0 | 8 | 0 | 0 | 0 | 0 | 0 | 0 | 0 | 0 | 0 | 0 | 0 | 0 | 0 |
| | *exad1-1* | NaCl (100mM) | 0 | 0 | 0 | 10 | 0 | 0 | 0 | 0 | 0 | 18 | 0 | 0 | 0 | 61 | 53 | 0 | 13 | 29 | 0 | 0 | 0 | 0 | 0 | 0 | 28 | 0 | 23 | 0 | 8 | 0 | 0 | 0 | 0 | 0 | 0 | 0 | 0 | 0 | 0 | 0 | 0 | 0 |
| | *exad1-3* | NaCl (100mM) | 0 | 0 | 0 | 9 | 0 | 0 | 0 | 0 | 0 | 18 | 0 | 0 | 0 | 59 | 54 | 0 | 13 | 31 | 0 | 0 | 0 | 0 | 0 | 0 | 32 | 0 | 27 | 0 | 9 | 0 | 0 | 0 | 0 | 0 | 0 | 0 | 0 | 0 | 0 | 0 | 0 | 0 |
| | *exad1-3* | NaCl (100mM) | 0 | 0 | 0 | 9 | 0 | 0 | 0 | 0 | 0 | 19 | 0 | 0 | 0 | 64 | 48 | 5 | 15 | 30 | 0 | 0 | 0 | 0 | 0 | 0 | 38 | 0 | 27 | 0 | 8 | 0 | 0 | 0 | 0 | 0 | 0 | 0 | 0 | 0 | 0 | 0 | 0 | 0 |
| | *exad1-3* | NaCl (100mM) | 0 | 0 | 0 | 10 | 0 | 0 | 0 | 0 | 0 | 17 | 0 | 0 | 0 | 40 | 34 | 6 | 18 | 30 | 0 | 0 | 0 | 0 | 0 | 0 | 30 | 0 | 22 | 0 | 9 | 0 | 0 | 0 | 0 | 0 | 0 | 0 | 0 | 0 | 0 | 0 | 0 | 0 |

**Figure S7. Heatmap of Comprehensive Microarray Polymer Profiling (CoMPP) spot signals.** 5-day-old seedlings of *exad1-1*, *exad1-3* and Col-0 were transferred to liquid medium containing NaCl (100 mM) or Control (0 mM) for 48 h treatment before harvesting. AIR samples were used extracted for CoMPP analysis. Genotypes, treatments and extraction conditions of sample are shown on the left. The corresponding monoclonal antibodies (mAbs) with specificity are shown on the top (Table S5). Signal values were correlated to color intensity and shown with three biological replicates.

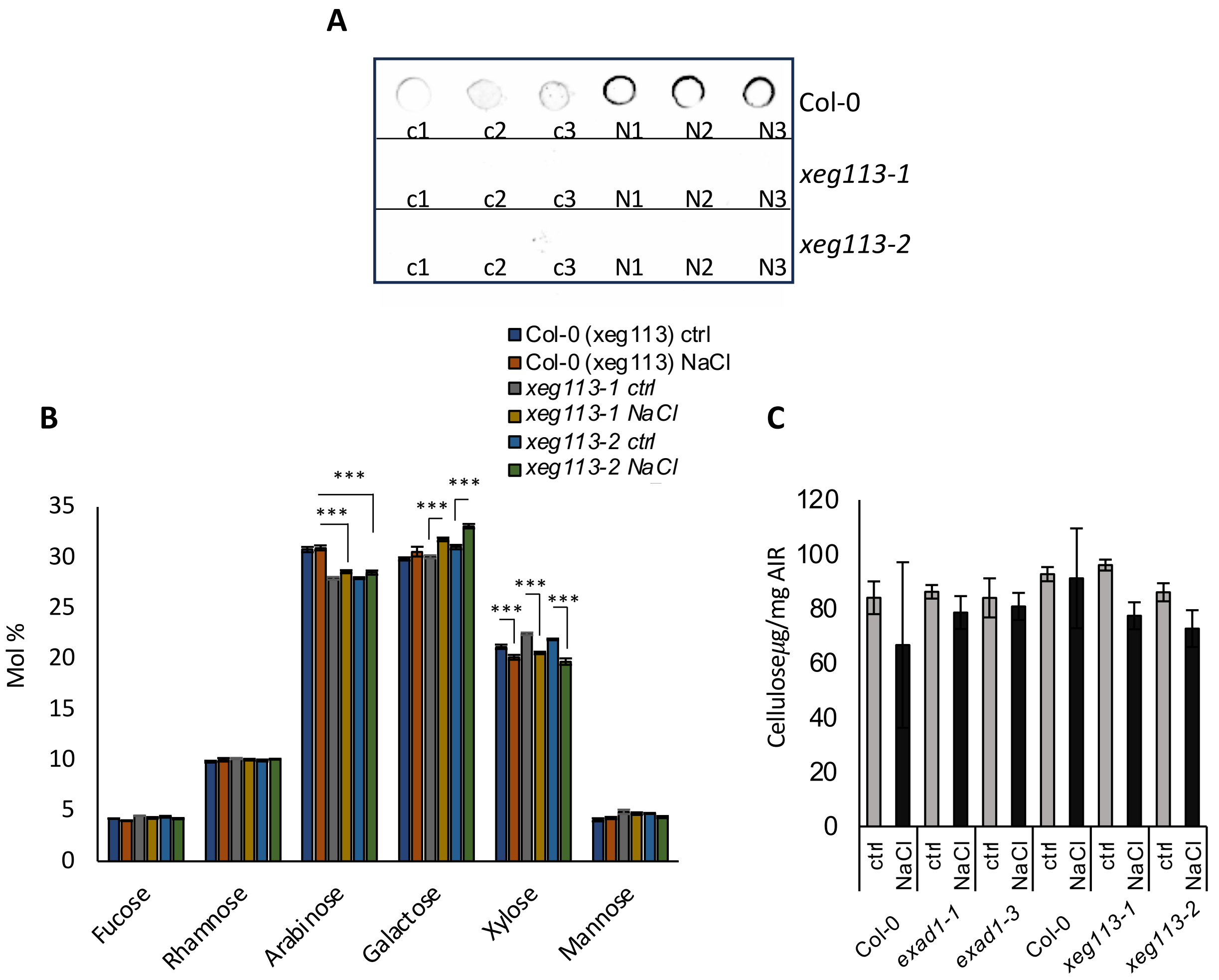


**Figure S8**. **A**, Five-day old seedlings of Col-0, *xeg113-1* and *xeg113-2* mutants were treated with or without 100 mM NaCl for 48 h before harvesting. Total protein were extracted and 25 ug of total protein spotted on nitrocellulose membrane. JIM11 antibody was used to detect the specific Hyp-Ara$_4$ signal. Dot-blot images are representative of 2 independent experiments performed each containing 3 biological replicates per treatment [Control (c1, c2, c3), or NaCl (N1, N2, N3)] per genotype. **C,** Five-day old seedlings of Col-0 *xeg113-1* and *xeg113-2* mutants were treated with 100 mM NaCl or without salt (Ctrl) for 48 h. Neutral cell wall matrix components were extracted from AIRs derived from 3 biological replicates per treatment/genotype and expressed as Molar Percentage (Mol%). Error bars represent SD of the mean values analyzed for each biological replicate (n= 3). Asterisks indicate statistically significant differences according to Student's t-test compared to Col-0 NaCl (for Arabinose) or compared to the corresponding controls (for Galactose and Xylose) (*, P< 0.05; **P< 0.01 ***P< 0.001) . **E,** Cellulose content expressed as μg of Glucose/ mg AIR was analyzed in five-day old seedlings of Col-0, *exad1-1, exad1-3,* Col-0, *xeg113-1* and *xeg113-2* mutants seedlings treated as in C. Statistical analysis performed by using two-way ANOVA with contrast post-hoc. No significant difference has been detected.

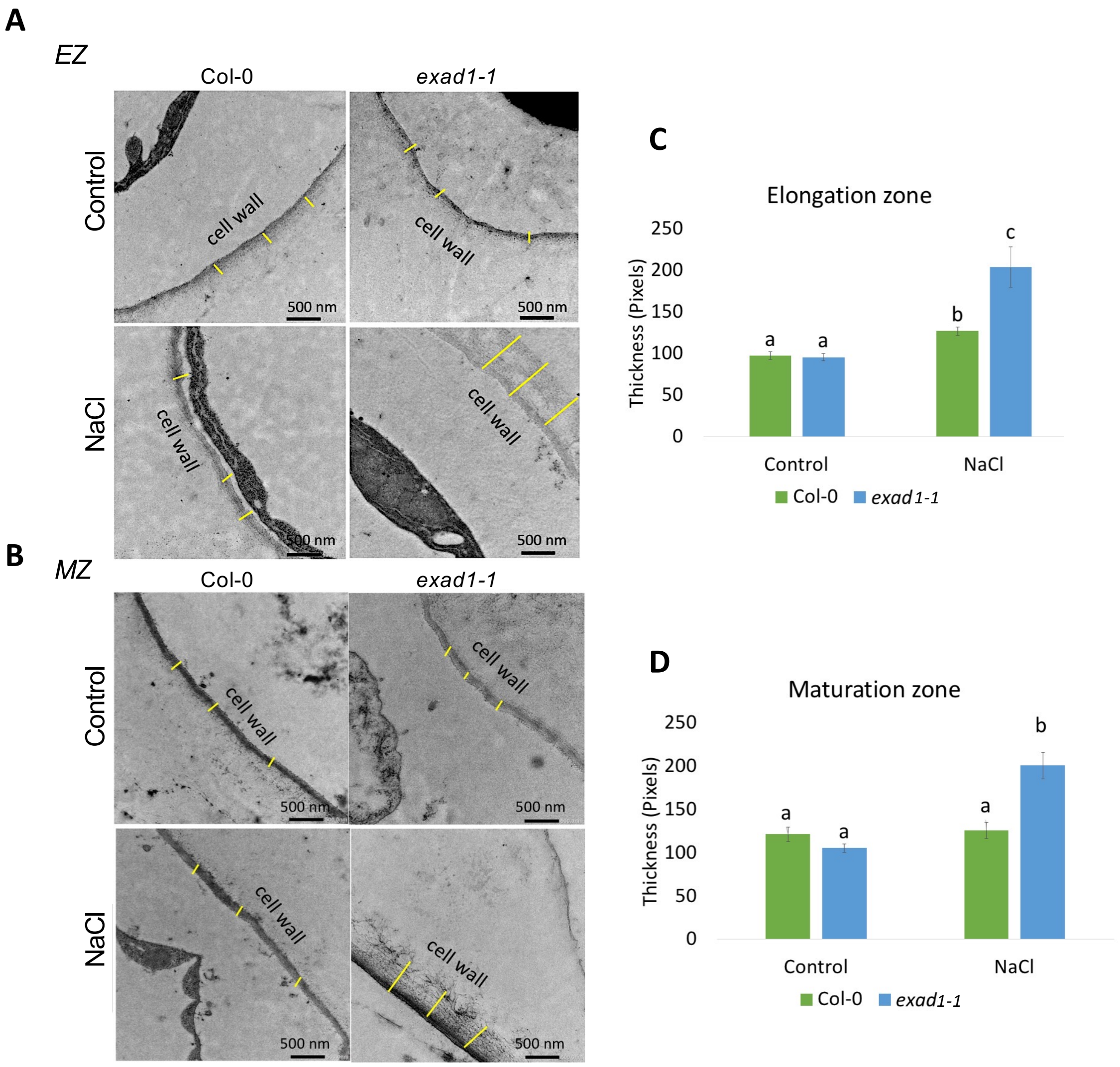


**Figure S9**. TEM analysis showing the cell wall thickness in epidermal cells of both the elongation zone (EZ) (**A**) and maturation zone (MZ) (**B**) of seedlings roots treated with and without NaCl (100 mM) for 48 h. Four-day old seedlings of Col-0 and *exad1-1* were transferred onto new plates in the SITA system. Samples were collected after 48 h treatments for TEM imaging. Cell walls and scale bar (= 500 nm) are indicated in the images. Quantifications of cell wall thickness in the elongation zone (**C**) and maturation zone (**D**) were measured from randomized selected 3 positions of cell wall (indicated as yellow lines) for each image. Statistical analysis was done using two-way ANOVA followed with contrasts post-hoc test, different letters indicate significant differences according to P-value < 0.05.
